# An ultraconserved snoRNA-like element in long noncoding RNA *CRNDE* promotes ribosome biogenesis and cell proliferation

**DOI:** 10.1101/2024.07.23.604857

**Authors:** Jong-Sun Lee, Tu Dan, He Zhang, Yujing Cheng, Frederick Rehfeld, James Brugarolas, Joshua T. Mendell

## Abstract

Cancer cells frequently upregulate ribosome production to support tumorigenesis. While small nucleolar RNAs (snoRNAs) are critical for ribosome biogenesis, the roles of other classes of noncoding RNAs in this process remain largely unknown. Here we performed CRISPRi screens to identify essential long noncoding RNAs (lncRNAs) in renal cell carcinoma (RCC) cells. This revealed that an alternatively-spliced isoform of lncRNA *Colorectal Neoplasia Differentially Expressed* containing an ultraconserved element (UCE), referred to as *CRNDE*^UCE^, is required for RCC cell proliferation. *CRNDE*^UCE^ localizes to the nucleolus and promotes 60S ribosomal subunit biogenesis. The UCE of *CRNDE* functions as an unprocessed C/D box snoRNA that directly interacts with ribosomal RNA precursors. This facilitates delivery of eIF6, a key 60S biogenesis factor, which binds to *CRNDE*^UCE^ through a sequence element adjacent to the UCE. These findings highlight the functional versatility of snoRNA sequences and expand the known mechanisms through which noncoding RNAs orchestrate ribosome biogenesis.

## INTRODUCTION

Uncontrolled cell growth and proliferation in cancer is fueled by high levels of translation, which necessitates an abundance of ribosomes^1^. To fulfill this heightened demand, cancer cells frequently exploit the intricate regulatory networks governing ribosome biogenesis^2,3^. For example, key drivers of growth and proliferation, such as the oncogenic transcription factor MYC and the mTOR pathway, stimulate ribosome biogenesis by increasing ribosomal RNA (rRNA) transcription, ribosomal protein translation, and ribosome assembly^4,5^. These pathways are highly active in many tumor types, including renal cell carcinoma (RCC), a deadly malignancy with the highest mortality rate among genitourinary cancers^6,7^. It has been recognized for more than a century that cancer cells frequently exhibit an increase in the size and abundance of nucleoli, the site of ribosome assembly^8,9^. This phenomenon is particularly striking in RCC, where nucleolar prominence is a key determinant of tumor grade^10^. These observations suggest that pathways that drive aberrant ribosome biogenesis may represent a therapeutic liability that can be exploited in RCC and other malignancies.

Ribosome biogenesis is a highly-regulated process that begins in the nucleolus with transcription of 47S precursor ribosomal RNA (pre-rRNA), which is processed to produce the mature 18S, 5.8S, and 28S rRNAs^11,12^. Coupled to each step of pre-rRNA processing is the sequential addition of ribosomal proteins, ultimately generating the 40S and 60S ribosomal subunits. This process is guided by a large suite of assembly factors (AFs) and closely monitored by multiple checkpoint and surveillance pathways that enable the tightly-controlled production of functional ribosomes^13^. The maturation of the 60S subunit in particular is a complex process involving nucleolar, nucleoplasmic, and cytoplasmic assembly stages^14^. A central player in 60S biogenesis is eukaryotic initiation factor 6 (eIF6), a highly conserved protein that is essential for 60S assembly across species^15^. In *Saccharomyces cerevisiae* and mammalian cells, eIF6 (known as TIF6 in yeast) associates with pre-60S subunits in the nucleolus, where it promotes rRNA processing and 60S subunit maturation before export of the subunit to the cytoplasm^16–18^. During the final stages of 60S biogenesis in the cytoplasm, eIF6 acts as a ribosome anti-association factor, preventing premature association of 60S subunits with 40S subunits^19–21^. Eviction of eIF6 from mature 60S subunits, which is required for the assembly of actively translating 80S ribosomes, is mediated by the Shwachman-Bodian-Diamond syndrome protein (SBDS) and the GTPase elongation factor-like 1 (EFL1)^22–24^. Following its release, eIF6 shuttles back to the nucleus, where it is recycled for further rounds of 60S subunit biogenesis. Underscoring the importance of this step of ribosome maturation, deficiency of SBDS or EFL1 causes Shwachman-Diamond Syndrome (SDS), a rare inherited ribosomopathy characterized by bone marrow failure, pancreatic insufficiency, skeletal abnormalities, and predisposition to hematopoietic malignancies^25–27^.

Noncoding RNAs also play key roles in ribosome biogenesis. In particular, small nucleolar RNAs (snoRNAs), a family of ∼60-250 nucleotide RNAs, serve as guides for site-specific chemical modification of rRNA and are critical for proper ribosome assembly and function^28^. snoRNAs can be classified into two major groups: C/D box snoRNAs, which guide 2’-*O*-methylation, and H/ACA box snoRNAs, which guide pseudouridylation. Each family of snoRNAs associates with a distinct set of co-factors to produce the functional snoRNA ribonucleoprotein particles (snoRNPs) that install the modifications in pre-rRNA. In addition to guiding nucleotide modifications, some snoRNAs promote pre-rRNA processing^29^, illustrating how base-pairing interactions between snoRNAs and rRNA orchestrate multiple aspects of ribosome biogenesis.

In metazoans, the vast majority of snoRNAs are encoded in introns of messenger RNAs (mRNAs) or noncoding RNAs^29,30^. After excision of the snoRNA-containing intron and debranching, exonucleolytic trimming from the 5’ and 3’ ends produces the mature snoRNA sequence, which is protected by association with snoRNP components. Interestingly, when two snoRNA sequences are present in a single intron, longer noncoding RNAs can be produced, with the tandem snoRNAs at each end protecting the intervening sequence from degradation^31^. One such noncoding RNA, termed *SLERT*, promotes pre-rRNA transcription^32^. Another lncRNA, known as *LoNA*, contains an unprocessed C/D box snoRNA-like element that negatively regulates 2’-*O*-methylation of pre-rRNA by sequestering snoRNP components^33^. These examples underscore the diversity of mechanisms through which snoRNA-like sequences can regulate ribosome biogenesis. Whether additional snoRNA-related RNAs, or other classes of noncoding RNAs, directly participate in later stages of ribosome assembly remains unknown.

In this study, we set out to identify long noncoding RNA (lncRNAs), a class of noncoding RNAs >200 nucleotides in length with diverse functions^34^, that are essential for growth of RCC cells. To this end, CRISPR interference (CRISPRi) screens were performed in multiple RCC cell lines, using a custom sgRNA library targeting approximately 750 lncRNAs that are overexpressed in this tumor type. These screens revealed that the lncRNA *Colorectal Neoplasia Differentially Expressed* (*CRNDE*) is essential for proliferation of all tested RCC cell lines. While *CRNDE* overexpression is linked to many cancers^35–38^, we discovered that a distinct isoform containing an ultraconserved element (UCE), termed *CRNDE*^UCE^, is essential in RCC cells. We further demonstrated that *CRNDE*^UCE^ localizes to the nucleolus and is required for the biogenesis of 60S ribosomal subunits. We found that the UCE in *CRNDE* functions as an unprocessed C/D box snoRNA-like element, associating with snoRNP components and directly interacting with 32S pre-rRNA. This interaction facilitates the delivery of eIF6, which binds to a sequence element adjacent to the UCE in *CRNDE*^UCE^, to pre-60S subunits, thus promoting ribosome biogenesis. Together, these findings illuminate the diverse mechanisms through which noncoding RNAs and snoRNA sequences impact ribosome biogenesis and illustrate how these functions are co-opted in cancer.

## RESULTS

### CRISPRi screening reveals that *CRNDE* is required for proliferation of RCC cells

Despite advances in targeted therapies and immunotherapies, RCC remains a deadly disease^39,40^, underscoring the need to identify new targets and pathways that drive this malignancy. Given their potential to function as oncogenic drivers^41^, we sought to systematically identify lncRNAs that are essential for proliferation and survival of RCC cells. To achieve this, we first generated a comprehensive catalog of lncRNAs that are highly expressed in this tumor type. RNA-seq data from a well-curated set of matched RCC tumor-normal pairs collected at UT Southwestern^42–44^, as well as RCC data from The Cancer Genome Atlas (TCGA)^45^, was analyzed to identify overexpressed lncRNAs (**Figures 1A** and **S1A**). These candidate lncRNAs were further filtered by selecting those that were detectably expressed in kidney cancer cell lines, as documented by the Cancer Cell Line Encyclopedia (CCLE)^46^. We also included 181 additional lncRNAs, representing the most highly expressed lncRNAs in RCC patient tumor samples not already selected. This yielded a final set of 753 genes, comprising the most overexpressed and abundant lncRNAs in RCC (**Table S1**).

**Figure 1.**
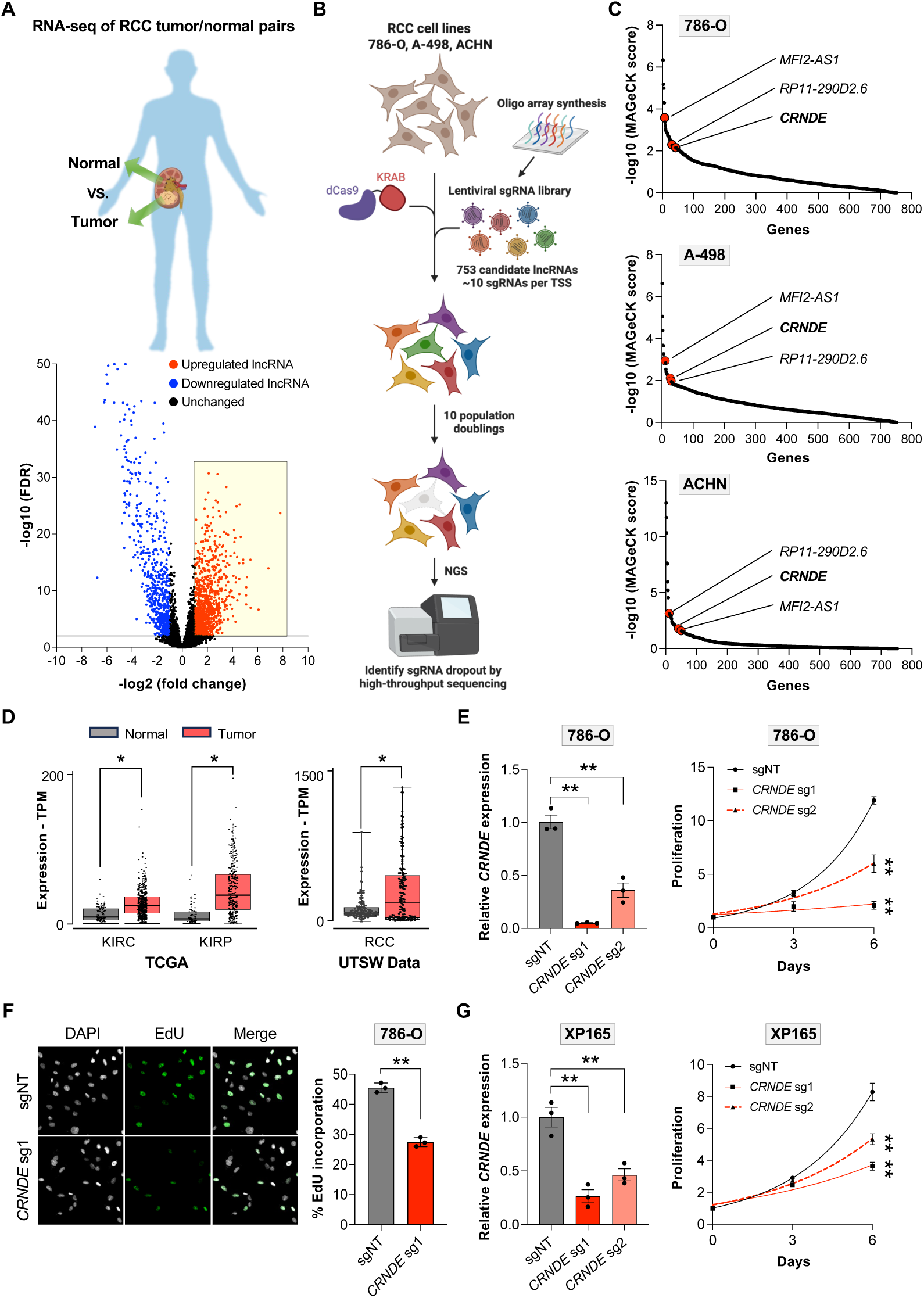
CRISPRi screening reveals that *CRNDE* is required for proliferation of RCC cells. (**A**) RNA-seq analysis of 48 matched tumor-normal pairs from RCC patients. The highlighted box indicates the lncRNAs whose expression level is at least 2-fold higher in tumors (FDR<0.05). (**B**) Schematic of CRISPRi drop-out screens performed in RCC cell lines. (**C**) MAGeCK analysis of CRISPRi screens. The common essential lncRNAs that were among the top 50 hits in all RCC cell lines are indicated. (**D**) TCGA (left) and UTSW RNA-seq data (right) showing elevated expression of *CRNDE* in RCC tumors. KIRC, Renal Clear Cell Carcinoma; KIRP, Renal Papillary Cell Carcinoma; RCC, all renal cell carcinoma subtypes. (**E**) qRT-PCR analysis of *CRNDE* expression relative to *GAPDH* in 786-O CRISPRi cells after lentiviral expression of non-target (sgNT) or *CRNDE*-targeting sgRNAs (left). Proliferation was measured by cell counts at the indicated timepoints after plating (right). (**F**) Fluorescence microscopy (left) and quantification (right) of DAPI and EdU incorporation in 786-O CRISPRi cells expressing the indicated sgRNAs. (**G**) qRT-PCR analysis of *CRNDE* expression relative to *GAPDH* in patient-derived primary RCC CRISPRi cells after lentiviral expression of non-target (sgNT) or *CRNDE*-targeting sgRNAs (left). Proliferation was measured by cell counts at the indicated timepoints after plating (right). Data are represented as mean ± SD (n=3 biological replicates). *p<0.05, **p<0.01, calculated by two-tailed t-test. **See also Figure S1.**

We next performed CRISPRi screens in multiple RCC cell lines (786-O, A-498, and ACHN), representing the most common RCC subtypes and molecular alterations^47^, to determine which of these lncRNAs are essential for RCC cell viability and/or proliferation. A lentiviral single guide RNA (sgRNA) library targeting the 753 candidate lncRNAs, consisting of approximately 10 sgRNAs per transcription start site (TSS), was constructed (**Figure 1B** and **Table S1**). sgRNAs targeting a set of known essential genes were included as positive controls. Stable cell lines expressing nuclease-deficient Cas9 fused to the KRAB transcriptional repressor domain (dCas9-KRAB) were generated, transduced with the library at high coverage (>1000 cells per sgRNA), and cultured for 10 population doublings. High-throughput sequencing was then performed and analyzed using MAGeCK^48^ to identify essential lncRNAs, representing those targeted by multiple sgRNAs that dropped out after prolonged passaging.

As expected, positive control sgRNAs targeting common essential genes were strongly depleted in all cell lines, demonstrating technical success of the screens (**Figure S1B**). Among the targeted lncRNAs, only three were shared among the top 50 most depleted lncRNAs in all three RCC cell lines: *MFI2-AS1*, *RP11-290D2.6*, and *CRNDE* (**Figure 1C** and **Table S2**). Both *MFI2-AS1* and *RP11-290D2.6* are antisense transcripts that fully overlap protein-coding genes, suggesting that silencing their expression is likely to impact the overlapping genes in *cis*. We therefore focused on *CRNDE* due to its potential to function through a *trans*-acting mechanism. *CRNDE* is a well-conserved intergenic lncRNA that has previously been shown to be overexpressed in many types of cancer, including RCC^35–38^. Indeed, analysis of TCGA and UT Southwestern RNA-seq data confirmed that *CRNDE* is strongly overexpressed in RCC (**Figure 1D**), and high expression is associated with poor patient survival (**Figure S1C**). These findings suggested an oncogenic role for *CRNDE* in RCC.

As predicted from the screening results and previous studies using short-hairpin-mediated knockdown^38^, depletion of *CRNDE* using CRISPRi strongly inhibited growth of all tested RCC cell lines (**Figures 1E, S1D,** and **S1E**). Incorporation of 5-ethynyl-2’-deoxyuridine (EdU) was reduced in *CRNDE*-deficient cells, indicative of reduced DNA replication (**Figure 1F**), while cleaved caspase-3, a marker of programmed cell death, remained undetectable (**Figure S1F**). Thus, loss of *CRNDE* impairs RCC cell proliferation without triggering apoptosis. We further showed that CRISPRi-mediated silencing of *CRNDE* impaired growth of multiple lines of primary patient-derived RCC cells (**Figures 1G** and **S1G**), demonstrating that the requirement for *CRNDE* is not limited to established RCC cell lines. Altogether, these results documented an essential role for *CRNDE* in RCC cell proliferation.

### A *CRNDE* isoform containing an ultraconserved element promotes RCC proliferation in *trans*

Although there is an extensive body of literature demonstrating that *CRNDE* knockdown impairs the proliferation and survival of many cancer cell lines^37^, no consensus exists regarding the molecular function of this lncRNA. LncRNAs may act by regulating local gene expression in *cis*, or they may leave the site of transcription and perform functions in *trans*^49,50^. The TSS of *CRNDE* is located ∼2 kilobases (kb) from the TSS of *Iroquois Homeobox 5* (*IRX5*) **(Figure S1H**), while no other genes are located within hundreds of kb of this locus. Importantly, CRISPRi-mediated knockdown of *CRNDE* did not affect expression of *IRX5* mRNA or protein (**Figures S1I** and **S1J**), arguing against local gene regulation by this lncRNA.

To test whether *CRNDE* functions in *trans*, we performed rescue experiments by expressing *CRNDE* in 786-O cells using a doxycycline (dox)-inducible PiggyBac transposon vector^51^. Expression of the full-length, unspliced *CRNDE* sequence rescued cell proliferation upon knockdown of endogenous *CRNDE*, demonstrating that a transcript produced from this locus supports RCC cell growth in *trans* (**Figures 2A** and **S2A**). Surprisingly, introduction of the fully-spliced *CRNDE* transcript was unable to rescue cell growth (**Figures 2A** and **S2B**), suggesting that this function might be attributable to an alternatively-spliced *CRNDE* isoform. Interestingly, analysis of ENCODE RNA-seq data revealed an abundance of reads spanning the final three introns of *CRNDE*, suggesting inefficient splicing (**Figure 2B**). Indeed, publicly-available long-read RNA sequencing data confirmed the expression of alternatively-spliced *CRNDE* isoforms containing these introns in multiple tissues (**Figure 2C**). Notably, these intron-containing transcripts harbored an ultraconserved element (UCE), a sequence of at least 200 nucleotides in length that exhibits extreme conservation among vertebrates^52,53^ (**Figure 2D**). Transcripts containing the alternatively-spliced introns were readily detectable by oligo dT-primed RT-PCR in 786-O cells (**Figure S2C**), while northern blotting using a probe complementary to the UCE confirmed the expression of a ∼5 kb *CRNDE* isoform containing this element (**Figure 2E**). Copy number analysis in RCC and other human cell lines documented the abundant expression of UCE-containing *CRNDE* transcripts in the range of ∼50-300 copies per cell (**Figure S2D**). Rescue experiments using the alternatively-spliced ∼5 kb *CRNDE* isoform containing the UCE demonstrated that expression of this transcript was sufficient to rescue proliferation upon knockdown of endogenous *CRNDE* (**Figure 2F**). Moreover, isolated expression of the third intron of *CRNDE* containing the UCE, or truncated versions with ∼800 nucleotides removed from the 5’ or 3’ ends, also supported cell proliferation (**Figures 2G, 2H, S2E,** and **S2F**; UCE intron, UCE intron 5’ deletion, or UCE intron 3’ deletion). Notably, the UCE alone or an upstream intron fragment lacking the UCE was not able to rescue growth (**Figures 2G, 2H** and **S2F**; UCE and UCE intron upstream). Together, these data demonstrated that an alternatively-spliced *CRNDE* isoform containing a UCE, hereafter referred to as *CRNDE*^UCE^, is required for RCC cell proliferation.

**Figure 2.**
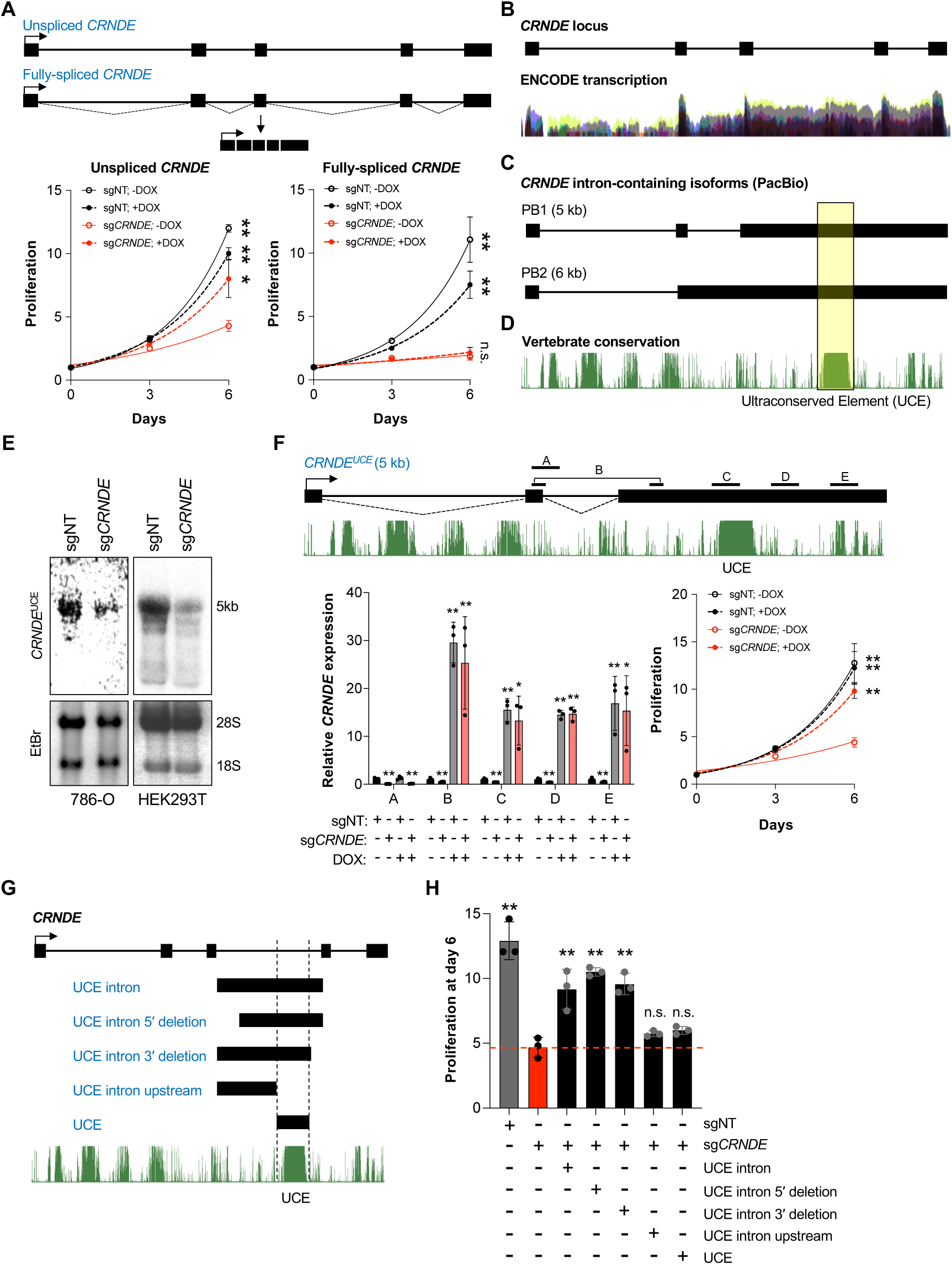
A *CRNDE* isoform containing an ultraconserved element promotes RCC proliferation in *trans*. (**A**) Proliferation of 786-O CRISPRi cells expressing doxycycline (dox)-inducible unspliced (left) or fully-spliced (right) *CRNDE* rescue constructs after lentiviral expression of non-target sgRNA (sgNT) or sgRNA targeting endogenous *CRNDE*. (**B-D**) UCSC genome browser ENCODE transcription (B) and vertebrate conservation (PhastCons) (D) tracks (hg38), aligned with intron-containing *CRNDE* isoforms detected in the PacBio Universal Human Reference RNA dataset^74^ (C). (**E**) Northern blot analysis of total RNA from the indicated cell lines after lentiviral expression of non-target (sgNT) or *CRNDE*-targeting sgRNAs using a probe complementary to the UCE. (**F**) qRT-PCR analysis (bottom left) and proliferation (bottom right) of 786-O CRISPRi cells expressing a dox-inducible *CRNDE* UCE-containing isoform rescue construct after lentiviral expression of non-target sgRNA (sgNT) or sgRNA targeting endogenous *CRNDE*. The positions of qRT-PCR amplicons (A, B, C, D, and E) are indicated in the transcript schematic. Transcript abundance was normalized to *GAPDH*. (**G-H**) Schematic showing positions of UCE intron rescue constructs (G) and their ability to support proliferation upon depletion of endogenous *CRNDE*, as measured by cell counts six days after plating (H). Data are represented as mean ± SD (n=3 biological replicates). n.s., not significant; **p<0.01, calculated by two-tailed t-test. **See also Figure S2.**

### *CRNDE*^UCE^ localizes to the nucleolus and promotes 60S ribosomal subunit biogenesis

To elucidate the molecular function of *CRNDE*^UCE^ underlying its requirement for cell proliferation, we next examined its subcellular localization. Cellular fractionation demonstrated that the fully-spliced *CRNDE* isoform localizes primarily to the cytoplasm in RCC cells, as reported in other cell types^54,55^ (**Figures 3A and S3A**). *CRNDE*^UCE^, in contrast, was highly enriched in nuclear fractions. Moreover, RNA fluorescence in situ hybridization (FISH) using probes complementary to the UCE revealed strong localization to the nucleolus (**Figure 3B**). To further validate nucleolar localization of *CRNDE*^UCE^, we performed pre-ribosome sequential extraction (PSE), which enables isolation of cytoplasmic and nucleoplasmic components (SN1 fraction), nucleolar ribosomes and associated components at late stages of maturation (SN2 fraction), and early pre-ribosomes near the core of the nucleolus (SN3 fraction) (**Figures 3C** and **3D**)^56,57^. Analysis of *CRNDE*^UCE^ abundance in these fractions verified its strong enrichment in the nucleolar interior (SN3) along with 32S pre-rRNA, a marker of early pre-ribosomes (**Figures 3E** and **S3B**).

**Figure 3.**
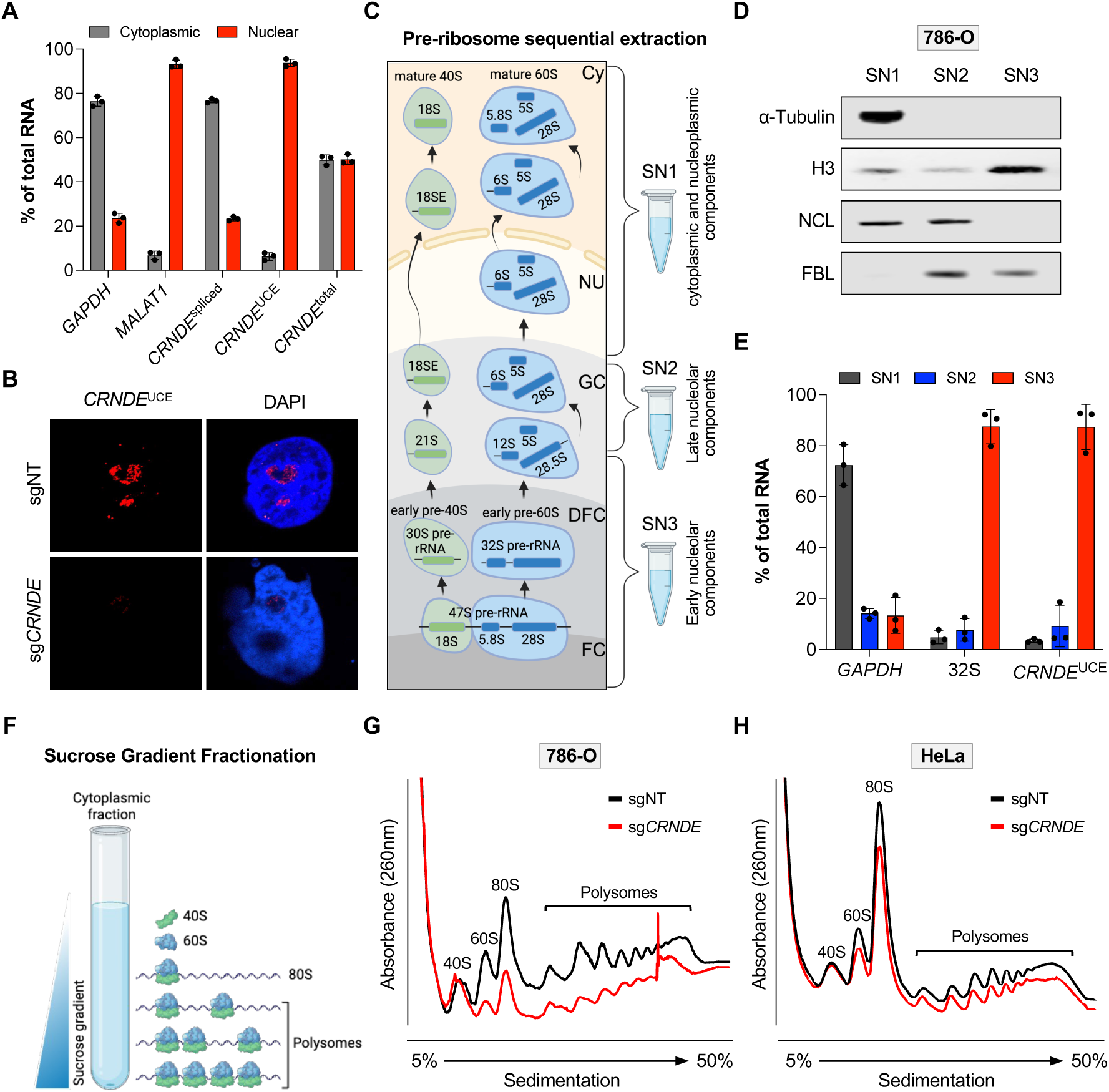
*CRNDE*^UCE^ localizes to the nucleolus and promotes 60S ribosomal subunit biogenesis. (**A**) Subcellular fractionation and qRT-PCR analysis of 786-O cells using primers that detect fully-spliced *CRNDE* (*CRNDE*^spliced^), *CRNDE* UCE-containing isoforms (*CRNDE*^UCE^), or all *CRNDE* isoforms (*CRNDE*^total^). *GAPDH* and *MALAT1* represent cytoplasmic and nuclear controls, respectively. (**B**) RNA FISH with a probe complementary to the UCE (red) in 786-O CRISPRi cells expressing the indicated sgRNAs. (**C**) Schematic of pre-ribosomal sequential extraction showing stages of ribosome biogenesis captured in each fraction (SN1, SN2, SN3). FC, fibrillar center; DFC, dense fibrillar component; GC, granular component; Nu, nucleoplasm; Cy, cytoplasm. (**D**) Western blot analysis of control proteins in nucleolar fractions. (**E**) qRT-PCR analysis of *CRNDE*^UCE^ in nucleolar fractions from 786-O cells. *GAPDH* and 32S pre-rRNA represent cytoplasmic and early nucleolar markers, respectively. (**F-H**) Schematic of polysome profiling by sucrose gradient ultracentrifugation (F) and results from 786-O (G) and HeLa (H) CRISPRi cells after lentiviral expression of non-target (sgNT) or *CRNDE*-targeting sgRNAs. Data are represented as mean ± SD (n=3 biological replicates). **See also Figure S3.**

Given that the nucleolus is the major site of ribosome biogenesis, we next examined the impact of loss of *CRNDE* on ribosomal subunit abundance and translation. Using sucrose density gradient ultracentrifugation, we observed that *CRNDE* deficiency resulted in a significant decrease in the levels of mature 60S subunits and a collapse of translating polysomes, while minimally affecting the abundance of 40S subunits (**Figures 3F, 3G** and **S3C**). To extend these analyses beyond RCC cells, we also examined *CRNDE* localization and function in HeLa cells and TERT-immortalized human fibroblasts (BJ). Depletion of *CRNDE* in these cell lines also impaired cell proliferation and measurably reduced 60S ribosomal subunit abundance, albeit to a lesser extent than in RCC cells (**Figures 3H** and **S3D-F**). Fractionation experiments demonstrated that *CRNDE*^UCE^ was similarly enriched within the nucleolar interior in these cells (**Figures S3G** and **S3H**). These data established that *CRNDE*^UCE^ is a nucleolar transcript that promotes the biogenesis of 60S ribosomal subunits in RCC and other cell types.

### *CRNDE*^UCE^ interacts directly with the 60S biogenesis factor eIF6

To determine how *CRNDE*^UCE^ promotes 60S ribosomal subunit biogenesis, we performed RNA antisense purification followed by mass spectrometry (RAP-MS) from UV-crosslinked RCC cells, which enables the identification of endogenous proteins that interact with a target RNA of interest^58^. To enhance the efficiency of detection of interacting proteins, we employed organic phase separation to enrich crosslinked RNPs prior to RNA pull-down^59,60^ (**Figures 4A** and **S4A**). Endogenous *CRNDE*^UCE^ RNP complexes were efficiently recovered using biotinylated antisense oligonucleotides (ASOs) complementary to the UCE region and co-purifying proteins were identified by mass spectrometry (**Figures S4B** and **S4C**). Notably, eIF6, a key factor in 60S ribosomal subunit biogenesis, was the most abundant protein uniquely detected in *CRNDE*^UCE^ pull-downs but not detected in control pull-downs. A direct interaction between *CRNDE*^UCE^ and eIF6 was confirmed by ASO-mediated pull-down of the crosslinked *CRNDE*^UCE^ RNP under denaturing conditions followed by western blotting (**Figure 4B**). Endogenous *CRNDE*^UCE^ was also highly enriched following reciprocal immunoprecipitation of endogenous eIF6 (**Figure 4C**).

**Figure 4.**
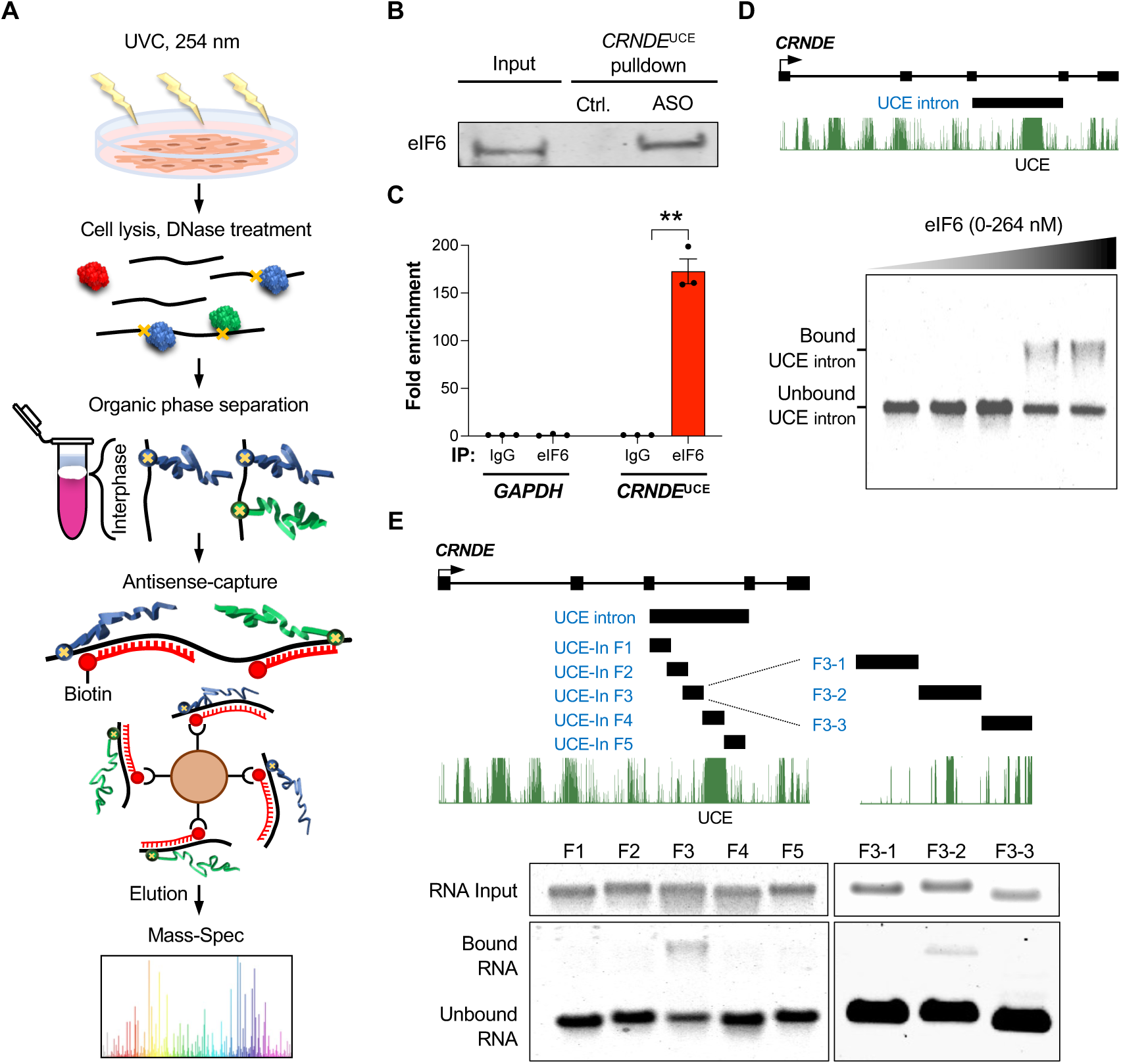
*CRNDE*^UCE^ interacts directly with the 60S biogenesis factor eIF6. (**A**) Schematic of UV-crosslinking and purification of endogenous *CRNDE*^UCE^ to identify interacting proteins by mass spectrometry. (**B**) Western blot of eIF6 after UV crosslinking and pull-down of *CRNDE*^UCE^ with ASOs under denaturing conditions. Scrambled ASO served as a negative control. (**C**) qRT-PCR analysis of *CRNDE*^UCE^ or *GAPDH* after UV-crosslinking and immunoprecipitation with anti-eIF6 antibody or control IgG. Fold enrichment over IgG plotted. Data are represented as mean ± SD (n=3 biological replicates). **p<0.01, calculated by two-tailed t-test. (**D-E**) Schematics of *CRNDE* UCE intron fragment (D, upper) or sub-fragments (E, upper) used in EMSAs with eIF6 (lower panels). **See also Figure S4.**

To further establish that eIF6 interacts directly with *CRNDE*^UCE^, electrophoretic mobility shift assays (EMSAs) were performed using in vitro transcribed UCE-containing intron fragments and purified Flag-tagged eIF6 (**Figure S4D**). eIF6 bound to the full-length UCE-containing intron (**Figure 4D**) as well as to a sub-fragment comprising the region immediately upstream of the UCE (**Figure 4E**; F3). We further narrowed the binding site to a 200-nucleotide sequence adjacent to the UCE (**Figure 4E**; F3-2). These results were consistent with our rescue experiments, which indicated that inclusion of the region upstream of the UCE was required to support growth of RCC cells (**Figures 2G, 2H** and **S2F**). These data raised the possibility that *CRNDE*^UCE^ promotes 60S biogenesis by regulating the activity of eIF6.

### *CRNDE*^UCE^ promotes loading of eIF6 onto pre-60S subunits in the nucleolus

Depletion of *CRNDE*^UCE^ had no effect on eIF6 mRNA or protein abundance (**Figures S5A** and **S5B**). Immunofluorescence, however, revealed a dramatic accumulation of eIF6 in nucleoli of *CRNDE*-depleted cells (**Figure 5A**). eIF6 is loaded onto pre-60S subunits in the nucleolus and it remains associated with 60S subunits until the final stages of maturation in the cytoplasm^15–21^. Therefore, the observed nucleolar accumulation of eIF6 in *CRNDE*-deficient cells suggested a defect in loading onto pre-ribosomes and/or export of pre-60S subunits to the cytoplasm.

**Figure 5.**
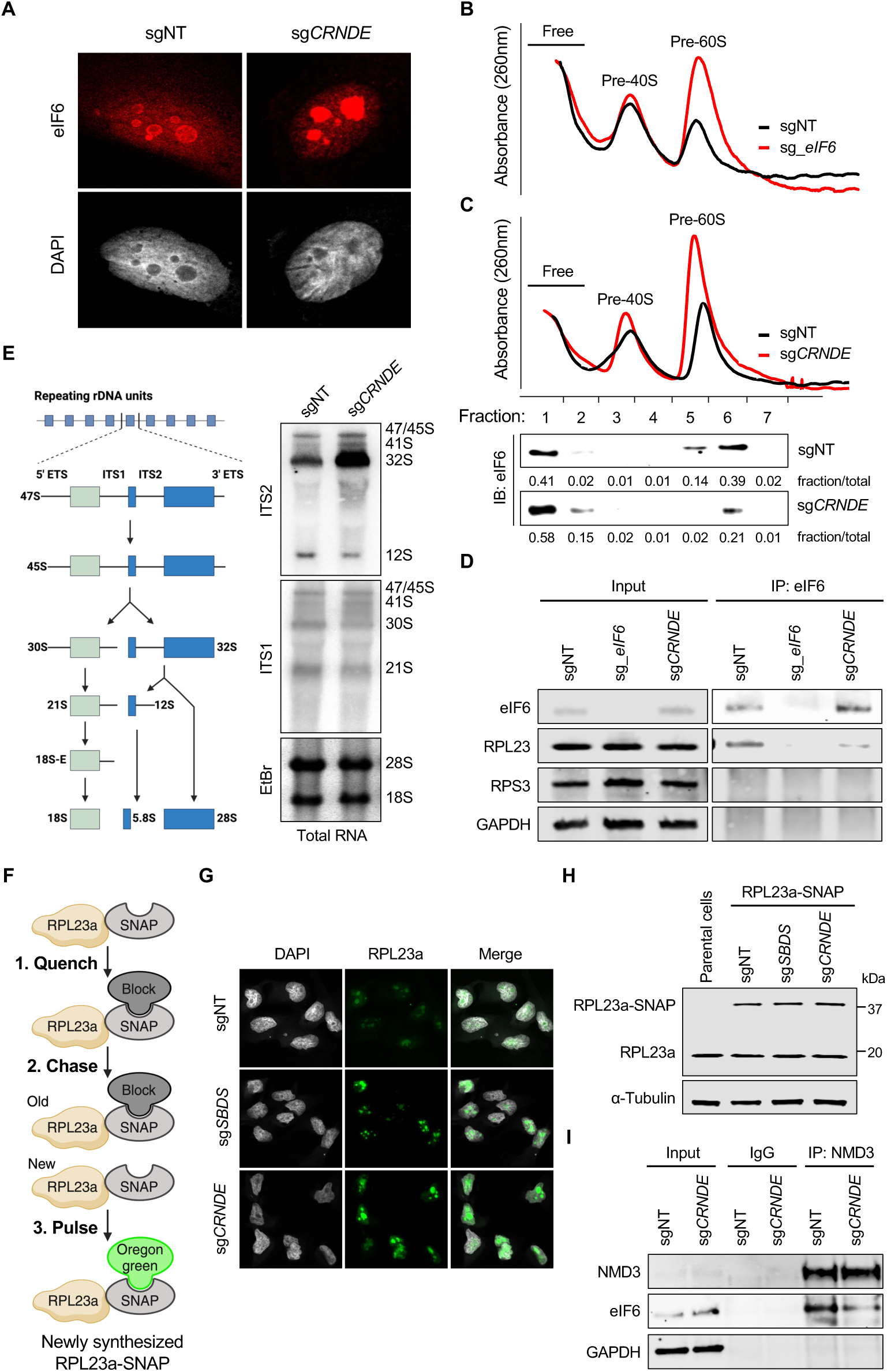
*CRNDE*^UCE^ promotes the loading of eIF6 onto pre-60S subunits in the nucleolus. (**A**) Immunolocalization of eIF6 in 786-O CRISPRi cells expressing the indicated sgRNAs. (**B-C**) Sucrose gradient fractionation of nucleoli from 786-O CRISPRi cells expressing sgRNAs targeting eIF6 (B) or *CRNDE* (C, upper). Western blot analysis of eIF6 in nucleolar fractions (C, lower). (**D**) Western blot analysis of input and eIF6 immunoprecipitates from 786-O CRISPRi cells expressing the indicated sgRNAs. (**E**) Schematic of pre-rRNA processing in human cells (left). Northern blot analysis of rRNA precursors with ITS1 and ITS2 probes in total RNA from 786-O cells (right). EtBr, ethidium bromide. (**F**) Schematic of SNAP labeling approach to detect newly synthesized large ribosomal subunits. (**G**) Fluorescence microscopy of newly-synthesized RPL23a-SNAP in 786-O CRISPRi cells expressing the indicated sgRNAs. (**H**) Western blot analysis of 786-O CRISPRi cells expressing RPL23a-SNAP and the indicated sgRNAs. (**I**) Western blot analysis of NMD3 or control IgG immunoprecipitates from 786-O CRISPRi cells expressing the indicated sgRNAs. **See also Figure S5.**

To examine eIF6 loading onto pre-60S subunits, we performed sucrose gradient fractionation of purified nucleoli. As expected, depletion of eIF6 led to a marked accumulation of pre-60S subunits in nucleoli, with only a minor impact on pre-40S subunit abundance (**Figure 5B**). Loss of *CRNDE* phenocopied these effects, providing additional evidence for an early block in large subunit maturation in cells deficient for this lncRNA (**Figure 5C**, upper). Western blotting further revealed a significant re-distribution of eIF6 from pre-60S fractions to ribosome-free fractions in *CRNDE* knockdown cells (**Figure 5C**, lower). Accordingly, immunoprecipitation demonstrated a reduced association of eIF6 with the large ribosomal subunit protein RPL23 upon depletion of *CRNDE* (**Figure 5D**). These data supported a role for *CRNDE*^UCE^ in promoting the loading of eIF6 onto pre-60S subunits.

eIF6 loading is coupled to subsequent pre-rRNA processing steps within the pre-60S subunit; specifically cleavage of 27S pre-rRNA in yeast and 32S pre-rRNA in human cells^17,61^. We therefore examined pre-rRNA processing intermediates in *CRNDE*-deficient cells using northern blotting. Consistent with the observed defect in eIF6 loading, we detected a strong accumulation of 32S pre-rRNA, along with a corresponding decrease in 12S pre-rRNA, in both total and nucleolar fractions (**Figures 5E** and **S5C**).

eIF6 loading is also required for the export of pre-60S subunits to the cytoplasm. To examine this step in ribosome biogenesis, we expressed a SNAP-tagged RPL23a protein that enabled specific labeling of newly-assembled pre-60S subunits whose trafficking from nucleoli could be monitored over time^62^ (**Figures 5F** and **5H**). After quenching pre-existing RPL23a-SNAP with a non-fluorescent benzyl-guanine substrate that reacts with the SNAP tag, newly-synthesized RPL23a-SNAP was labeled with Oregon Green and tracked by fluorescence microscopy. Loss of *CRNDE* resulted in a robust accumulation of newly-synthesized RPL23a-SNAP within nucleoli to an extent comparable to that observed upon depletion of SBDS, whose loss traps eIF6 in the cytoplasm and is known to result in a pre-60S export defect^23^ (**Figure 5G**). Furthermore, immunoprecipitation of the pre-60S export factor NMD3 revealed a reduced association with eIF6 upon *CRNDE* depletion, suggesting impaired formation of pre-60S export complexes (**Figure 5I**). Altogether, these data strongly supported a role for *CRNDE*^UCE^ in facilitating the loading of eIF6 onto maturing pre-60S subunits in the nucleolus, resulting in a defect in subsequent steps in ribosome biogenesis upon loss of this lncRNA.

### The *CRNDE* UCE is a C/D box snoRNA-like sequence that directly interacts with 32S pre-rRNA

We next investigated how the interaction of eIF6 with *CRNDE*^UCE^ promotes its loading onto pre-60S subunits. Given the key roles of snoRNAs in ribosome biogenesis, which base-pair with specific sites in pre-rRNA to guide nucleotide modifications and processing^28^, we hypothesized that *CRNDE*^UCE^ may similarly base-pair with pre-rRNA in order to facilitate eIF6 delivery to maturing pre-ribosomes. To examine this possibility, 786-O cells were treated with 4’-aminomethyltrioxalen (AMT), a psoralen-derived crosslinker that induces interstrand crosslinks between base-paired RNAs upon exposure to UV light^63^. *CRNDE*^UCE^ complexes were then selectively captured using biotinylated ASOs complementary to the UCE, subjected to stringent washing in urea-containing denaturing buffers^64^, and analyzed by qRT-PCR (**Figure 6A**). Remarkably, a robust enrichment of 32S pre-rRNA was detected in *CRNDE*^UCE^ pull-down samples, while mature 18S or 28S rRNA, or an unrelated mRNA (*GAPDH*), exhibited minimal enrichment (**Figure 6B**). Use of a scrambled ASO, or pull-downs performed in the absence of AMT crosslinking (**Figure S5D**), yielded no enrichment, further demonstrating the specificity of the 32S pre-rRNA signal in *CRNDE*^UCE^ purifications. Additionally, the *CRNDE*^UCE^:32S pre-rRNA interaction occurred independently of eIF6, as demonstrated by pull-down assays performed following eIF6 depletion by CRISPRi (**Figure 6C**). These results supported a direct base-pairing interaction between *CRNDE*^UCE^ and 32S pre-rRNA.

**Figure 6.**
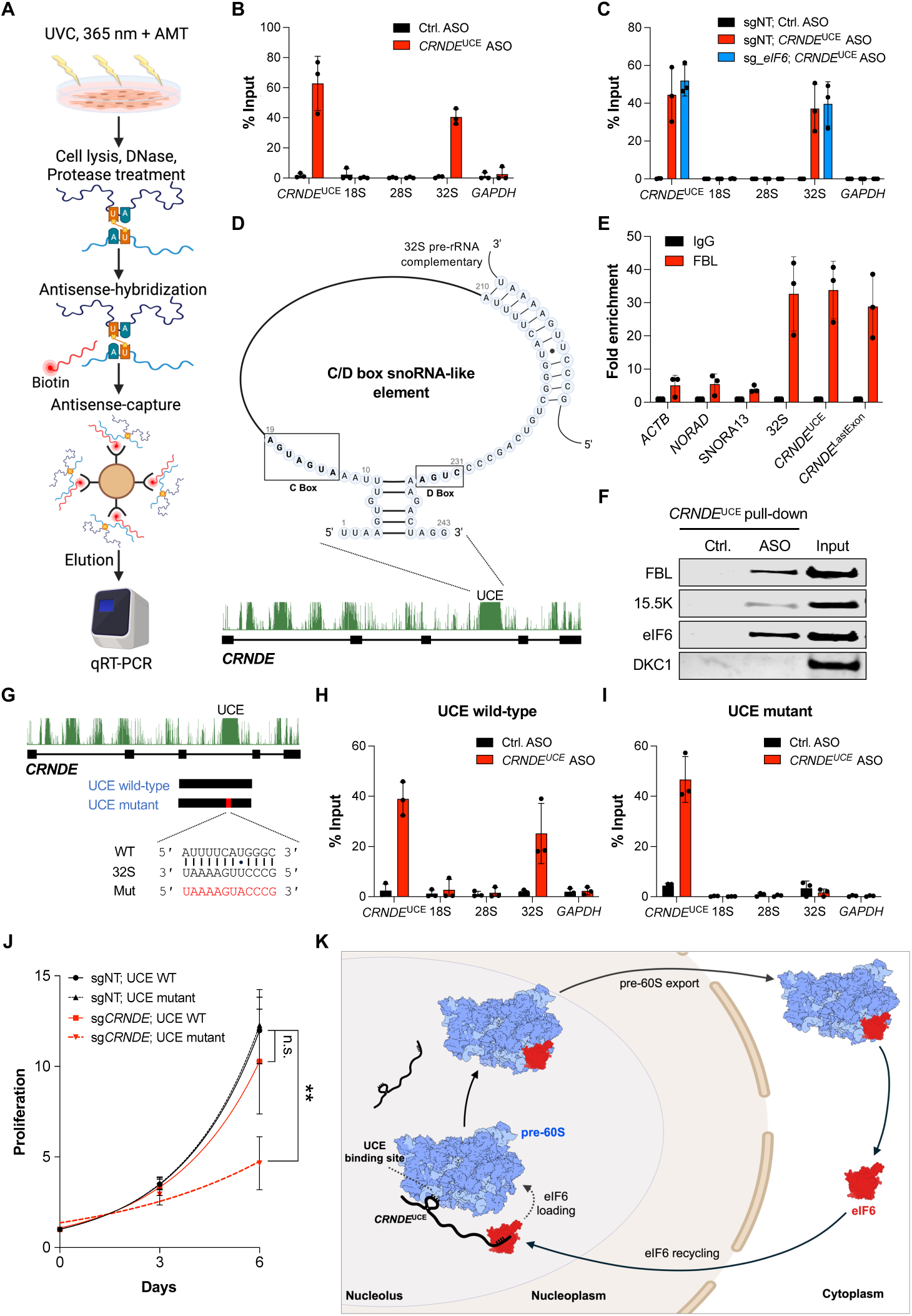
The *CRNDE* UCE is a C/D box snoRNA-like element that base pairs with 32S pre-rRNA. (**A**) Schematic of AMT crosslinking experiment to detect RNA:RNA base-pairing interactions. (**B-C**) qRT-PCR analysis of the indicated transcripts after AMT crosslinking and pull-down of *CRNDE*^UCE^ with ASOs under denaturing conditions. Experiments were performed in parental 786-O CRISPRi cells (B) or in 786-O CRISPRi cells expressing non-target (sgNT) or eIF6-targeting sgRNAs (C). Scrambled ASO served as a negative control. Enrichment was normalized to input. (**D**) The UCE of *CRNDE* resembles a C/D box snoRNA. Putative C box, D box, and complementarity to 32S pre-rRNA indicated. (**E**) qRT-PCR analysis of negative control transcripts (*ACTB*, *NORAD*, H/ACA box snoRNA SNORA13), 32S pre-rRNA (positive control), or *CRNDE* transcripts (*CRNDE*^UCE^ or *CRNDE*^LastExon^) in FBL immunoprecipitates from UV-crosslinked 786-O cells. Fold enrichment over IgG plotted. (**F**) Western blot analysis of C/D box snoRNP components FBL and 15.5K, or H/ACA box snoRNP component DKC1, after UV crosslinking and pull-down of *CRNDE*^UCE^ with ASOs under denaturing conditions. Scrambled ASO served as a negative control. (**G**) Schematic of *CRNDE* UCE intron fragments and sequence of 32S binding site mutant used in rescue experiments. (**H-I**) qRT-PCR analysis of the indicated transcripts after AMT crosslinking and pull-down of *CRNDE*^UCE^ from cells expressing wild-type (H) or mutant (I) UCE-containing intron fragments after depletion of endogenous *CRNDE* using CRISPRi. (**J**) Proliferation of 786-O CRISPRi cells expressing wild-type or mutant UCE-containing intron fragments after lentiviral expression of non-target sgRNA (sgNT) or sgRNA targeting endogenous *CRNDE*. (**K**) Regulation of 60S ribosomal subunit biogenesis by *CRNDE*^UCE^. The UCE functions as an unprocessed C/D box snoRNA that base-pairs with 32S pre-rRNA. This facilitates delivery of eIF6, bound to an adjacent sequence element, to maturing pre-60S subunits, thereby promoting subsequent steps in 60S biogenesis. Data are represented as mean ± SD (n=3 biological replicates). n.s., not significant; **p<0.01, calculated by two-tailed t-test. **See also Figure S5 and S6.**

We next sought to identify the region of *CRNDE*^UCE^ that base-pairs with 32S pre-rRNA. Intriguingly, we noticed that the *CRNDE* locus exhibits multiple blocks of intronic sequence conservation, a feature reminiscent of snoRNA host transcripts^29,30^ (**Figures 6D and S5E**). The UCE region in particular appeared to resemble a C/D box snoRNA, with sequences matching potential C box (AUGAUGA) and D box (CUGA) motifs adjacent to a potential six base-pair terminal stem. The guide sequences of C/D box snoRNAs, which base-pair directly with pre-rRNA, are typically located ∼10-21 nucleotides upstream of the D box. Consistent with this configuration, we noted the presence of a sequence with complementarity to 32S pre-rRNA within this window upstream of the *CRNDE*^UCE^ D box (**Figures 6D and S5F**). This 12-nucleotide putative guide sequence contains a single mismatch at the position at which the 2’-*O*-methyl modification would be installed by a canonical C/D box snoRNP. Accordingly, methylation at this position, which is located in the linker between helices H55 and H56 in 28S rRNA domain III (**Figures S6A and S6B**), has not been detected in prior studies of rRNA nucleotide modifications^65^.

To determine if the UCE of *CRNDE* interacts with components of C/D box snoRNPs, we performed UV-crosslinking and immunoprecipitation of endogenous fibrillarin (FBL), the methyltransferase guided by C/D box snoRNAs^66^ (**Figures 6E and S5G**). We observed strong enrichment of *CRNDE*^UCE^ and, as expected, 32S pre-rRNA, but not control RNAs, in FBL pull-downs. The specificity of this interaction was further confirmed by reciprocal pull-down of the endogenous *CRNDE*^UCE^ RNP from UV crosslinked cells under denaturing conditions using ASOs (**Figure 6F**). Both FBL and 15.5K, another C/D box snoRNP component, co-purified with *CRNDE*^UCE^, while Dyskerin (DKC1), a component of H/ACA box snoRNPs, was not detected. These data provide strong evidence that the UCE of *CRNDE* assembles a snoRNP-like complex. Nevertheless, we were not able to detect any evidence of processing of the UCE into a snoRNA-like small RNA. Northern blotting using a probe complementary to the UCE failed to detect any small RNA species (**Figure S5H**). Moreover, immunoprecipitation of FBL led to equal enrichment of the UCE and final exon of *CRNDE* (**Figure 6E**), suggesting that FBL associates with the unprocessed *CRNDE*^UCE^ isoform.

We next asked whether the putative guide sequence in the *CRNDE*^UCE^ snoRNA-like element mediates the observed interaction with 32S pre-rRNA. The wild-type UCE-containing intron sequence (UCE wild-type), or a version with a mutation in the putative guide sequence (UCE mutant), were expressed in 786-O cells, while endogenous *CRNDE* was silenced using CRISPRi (**Figures 6G** and **S5I**). AMT crosslinking and pull-down of the *CRNDE* UCE using ASOs revealed the expected enrichment of 32S pre-rRNA in cells expressing the wild-type UCE intron but not in cells expressing the mutant sequence, demonstrating that this region is required for interaction with pre-rRNA (**Figures 6H** and **6I**). Additionally, we observed that the UCE guide sequence mutation also abolished the ability of the UCE intron to support RCC cell growth (**Figure 6J**). In sum, these data demonstrated that the UCE of *CRNDE* functions as an unprocessed C/D box snoRNA-like sequence that interacts directly with 32S pre-rRNA. This interaction likely facilitates efficient loading of eIF6, which is bound to a sequence adjacent to this element, onto pre-60S subunits.

## DISCUSSION

Many prior studies have documented a role for *CRNDE* in cancer cell proliferation and survival, making it one of the most highly studied lncRNAs in human malignancies^35–38^. Nevertheless, no consensus exists regarding the molecular function of this lncRNA that underlies its hyperactivity in cancer. Here we describe a key role for *CRNDE* in ribosome biogenesis and demonstrate that this function is essential for proliferation of RCC cells. These findings deepen our understanding of how noncoding RNAs govern ribosome assembly and illuminate how cancer cells can co-opt these mechanisms to fuel tumor growth.

To identify lncRNAs that are essential for RCC cell proliferation and/or survival, we generated a custom CRISPRi library targeting lncRNAs that are overexpressed in RCC tumors. Screens in multiple RCC cell lines, followed by validation studies in patient-derived primary RCC cultures, revealed that *CRNDE* is required for growth of all tested RCC cells. Rescue experiments demonstrated that this activity is attributable to an alternatively-spliced isoform of *CRNDE* containing an ultraconserved element (*CRNDE*^UCE^). We found that this transcript localizes to the nucleolus where an unprocessed C/D box-snoRNA-like element within the UCE base-pairs with maturing rRNA precursors. *CRNDE*^UCE^ also interacts directly with eIF6, a critical factor for 60S ribosomal subunit biogenesis, and promotes loading of eIF6 onto pre-60S subunits and subsequent steps in pre-60S maturation. Based on these findings, we propose that *CRNDE*^UCE^ functions as a unique ribosome assembly factor that exploits both RNA:RNA and RNA:protein interactions to position eIF6 in close proximity to its binding site on the pre-60S subunit, thereby enhancing ribosomal subunit maturation (**Figure 6K**). Consistent with this model, the structure of a human nucleolar pre-60S particle containing eIF6^67^ shows that the *CRNDE*^UCE^ binding site in 32S pre-rRNA is on the surface of the particle in a structurally unresolved loop where it would likely be accessible for base-pairing (**Figure S6C**). This site is on the same face and within 120Å of the eIF6 binding site on the maturing ribosomal subunit, a distance that could easily be bridged by the ∼600 nucleotides separating the snoRNA-like element and the eIF6 binding site in *CRNDE*^UCE^. We posit, therefore, that base-pairing between *CRNDE*^UCE^ and 32S pre-rRNA at this site facilitates recruitment of the *CRNDE*^UCE^:eIF6 complex to the pre-60S particle, although it is also possible that *CRNDE*^UCE^ binding induces conformational changes in the pre-ribosomal subunit that promote eIF6 loading.

A significant challenge for understanding the functions of lncRNAs is the paucity of well-established functional RNA domains, whose presence in a transcript is predictive of a specific molecular activity. Our data, together with previously reported results, demonstrate that snoRNA-like sequences are a repeatedly used class of RNA elements that impact ribosome biogenesis in multiple ways. Classically, C/D box and H/ACA box snoRNAs function as guides for 2’-*O*-methylation and pseudouridylation of pre-rRNA, respectively. As such, they are strongly localized to the nucleolus and evolutionarily optimized to base-pair with pre-rRNA. These attributes can be exploited for regulation of other steps in ribosome production. For example, some snoRNAs direct specific pre-rRNA processing events^29^ or regulate the rate of incorporation of ribosomal proteins into maturing ribosomal subunits^68^. LncRNAs harboring partially processed or unprocessed snoRNA-like elements have been shown to regulate transcription and chemical modification of pre-rRNA^32,33^. Our discovery that *CRNDE*^UCE^ uses an unprocessed snoRNA-like element to interact with pre-rRNA in order to deliver a key assembly factor to maturing pre-60S subunits expands the known mechanisms through which this versatile class of RNA elements can impact ribosome assembly. Moreover, these results illustrate how the presence of a snoRNA-like sequence within a lncRNA should be regarded as a strong clue that the transcript may localize to the nucleolus and regulate ribosome production.

Many alternatively-spliced variants of *CRNDE* have been detected ^35,54^, providing a regulatable mechanism for controlling the activity of this lncRNA. Retention of intron 3 yields the *CRNDE*^UCE^ transcript containing the eIF6 binding site and the snoRNA-like element, which promote maturation of the pre-60S subunit. Splicing of this intron produces transcript variants that localize to the cytoplasm and may perform other cellular functions. Thus, signaling pathways and factors that regulate this splicing event may impact ribosome biogenesis. Notably, the location of a snoRNA sequence within its host intron is a critical determinant of the efficiency of its processing into a mature small RNA. The optimal location for an efficiently processed snoRNA is 70-80 nucleotides upstream of the 3’ splice site, while deviations from this optimum can impair snoRNA generation^69,70^. The snoRNA-like element in *CRNDE*^UCE^ is located approximately 1 kb upstream of the nearest 3’ splice site, providing a likely explanation for its failure to be processed into a mature snoRNA. This enables the physical coupling of the snoRNA-like sequence with the eIF6 binding element, allowing *CRNDE*^UCE^ to function as a bi-partite ribosome assembly factor.

In addition to our finding that *CRNDE* is overexpressed and essential for proliferation in RCC, this lncRNA has been demonstrated to support proliferation in many types of cancer cells^37,38,55,71,72^. This suggests that *CRNDE*^UCE^ may play a broad role in driving ribosome biogenesis in diverse human malignancies, although it is important to note that *CRNDE* has been reported to promote tumorigenesis through a variety of additional mechanisms such as modulating the activity of the Wnt/β-catenin, mitogen-activated protein kinase (MAPK), and PI3K/AKT/mTOR pathways^37^. In light of the extreme evolutionary conservation of the UCE, and our discovery of its role in ribosome biogenesis, it is surprising that complete deletion of the *Crnde* locus is compatible with overtly normal development in mice^73^. This suggests that, rather than functioning as a core component of the 60S biogenesis pathway, *CRNDE*^UCE^ may act as an enhancer of ribosome assembly that is selectively deployed in settings of increased ribosomal demand, such as in regenerating tissues or in response to other stresses that require high levels of protein synthesis. Given that *CRNDE* appears to be dispensable under normal physiologic conditions, but is essential in RCC and other malignancies, targeting this lncRNA, or the pathways that regulate it, may provide a well-tolerated therapeutic strategy for cancer.

Achieving this goal will require a deeper understanding of the physiologic roles of *CRNDE*^UCE^-regulated ribosome biogenesis, a priority for future study.

## ACKNOWLEDGEMENTS

We thank the patients who provided tumors analyzed in this study. We also acknowledge Eric Campeau, Nancy Craig, Paul Kaufman, Xiaojun Lian, Didier Trono, Rudolf Jaenisch, Jonathan Weissman, and Feng Zhang for plasmids; Vanessa Schmid in the McDermott Center Next Generation Sequencing Core for assistance with high-throughput sequencing; Andrew Lemoff in the UTSW Proteomics Core for assistance with mass spectrometry; Salman Banani for assistance with CRISPRi library design; Jan Erzberger for assistance with structural analyses of rRNA and ribosomes; and Michael Buszczak, Jan Erzberger, and members of the Mendell laboratory for helpful suggestions on the manuscript. This study utilized data generated by the TCGA Research Network: https://www.cancer.gov/tcga. This work was supported by grants from CPRIT (RP220309 to J.T.M.), the Welch Foundation (I-1961 to J.T.M.), DOD (W81XWH2110815 to J-S.L.), and NIH (R01CA282036 to J.T.M. and P50CA196516 to J.B.). J.T.M. is an Investigator of the Howard Hughes Medical Institute.

## AUTHOR CONTRIBUTIONS

J-S.L., D.T., Y.C., F.R., J.B., and J.T.M. designed experiments and interpreted the results. J-S.L., D.T., and Y.C. performed experiments. He Z. performed bioinformatic analyses. F.R. and J.B. provided technical assistance and critical reagents. J-S.L. and J.T.M. wrote the manuscript.

## MATERIALS AND METHODS

### Cell culture

786-O cells were cultured in RPMI (Invitrogen) supplemented with 10% fetal bovine serum (Sigma-Aldrich) and 1X Antibiotic-Antimycotic (Gibco). A-498 and ACHN cells were cultured in Minimum Essential Medium Eagle (Sigma-Aldrich) supplemented with 10% fetal bovine serum and 1X Antibiotic-Antimycotic. Primary patient-derived RCC cells were cultured in Minimum Essential Medium (Gibco) supplemented with 1X MEM non-essential amino acids (Gibco), Hydrocortisone (0.4 mg/mL, StemCellTechnologies), EGF (5 µg/mL, Gibco), 10% fetal bovine serum, and 1X Antibiotic-Antimycotic. HEK293T and HeLa cells were cultured in Dulbecco’s Modified Eagle’s Medium (DMEM) (Invitrogen) supplemented with 10% fetal bovine serum and 1X Antibiotic-Antimycotic. BJ cells were cultured in DMEM supplemented with 20% Medium 199 (Gibco), 10% fetal bovine serum, and 1X Antibiotic-Antimycotic. 786-O, A-498, ACHN, HEK293T, and HeLa cells were obtained from ATCC. Primary patient-derived RCC cells were previously described^44^. Cell lines were confirmed to be free of mycoplasma contamination.

### Generation of CRISPRi cell lines

RCC cell lines (786-O, A-498 and ACHN), HeLa cells, and BJ cells were infected with a dCas9^KRAB^-mCherry lentivirus (Addgene #60954)^77^ and clonal lines stably expressing high levels of mCherry were established by single-cell sorting and expansion. Primary patient-derived RCC CRISPRi cells (XP127 and XP165) were generated by infecting with dCas9^KRAB^-mCherry lentivirus and sorting the cells with high mCherry expression (top 30%).

### sgRNA library design

The UT Southwestern (UTSW) Kidney Cancer Program (KCP) platform was used to analyze primary patient RNA-seq data from 48 RCC tumors and their matched normal kidney samples obtained from UTSW-affiliated hospitals, including Parkland Hospital and Children’s Medical Center^44,78^. Of a total of 63,677 expressed transcripts in this dataset, 13,283 were assigned a noncoding RNA biotype (antisense, sense intronic, sense overlapping, TEC, and lincRNA). Transcripts with these biotypes that exhibited >2-fold upregulation in tumor samples compared to matched normal pairs, an expression value of ≥1 TPM in tumors, detectable expression in CCLE kidney cancer cell lines (786-O, A-498, Caki-2, and ACHN), and overexpression in TCGA RCC data were selected for sgRNA design (572 genes). Additionally, the 181 most highly expressed lncRNAs in UTSW RCC tumors that were not already selected were included. This yielded a final set of 753 candidate lncRNAs. sgRNAs were designed using a previously-described algorithm that incorporates information on chromatin state, guide position relative to transcription start site (TSS), predicted off-target sites, and other sequence features^79^. 10 sgRNAs per TSS were designed. 450 previously-designed non-targeting controls^80^ and 120 previously-designed depletion controls targeting common essential genes (PCNA, RPA1, RPA3, RPS15A, RPS19, RPS11, RPS21, RPL32, RPS18, RPL7A, RPL8, RPL6)^79^ were included in the library, producing a final library size of 11,814 sgRNAs. Oligonucleotides were synthesized by CustomArray and shotgun cloned into pU6-sgRNA lentiviral vector (Addgene #60955) as previously described^77^. Noncoding RNA targets and sequences of sgRNAs are provided in **Table S1**.

### CRISPRi screens in RCC cells

CRISPRi screens were performed in replicate using two independent dCas9^KRAB^ expressing clones per cell line. For each replicate, 5 × 10^7^ cells were seeded into fourteen 15 cm dishes, with each dish containing medium supplemented with 6 µg/mL of polybrene (Millipore) and the lentiviral library at a multiplicity of infection of ∼0.3. After overnight incubation, media was changed and cells were cultured for 2 days, followed by selection in 2 µg/mL puromycin for 3 days. Following selection, at least 1.2 × 10^7^ cells were plated for each replicate, representing ∼1000x coverage of the library. This timepoint was designated “day 0”. Cells were maintained in 1 µg/mL puromycin for 15 days, with passaging every 3 days, each time plating at least 1.2 × 10^7^ cells to maintain a minimum of 1000x library coverage. Cells were harvested at day 0 and day 15 for genomic DNA isolation. Genomic DNA was extracted using the MasterPure Complete DNA Purification kit (Lucigen). Sequencing libraries were generated through two sequential rounds of PCR using Herculase II Fusion DNA polymerase (Agilent). For the first round of PCR, 6.6 µg of genomic DNA (representing ∼10^6^ cells) was used in each 100 µL reaction. A total of 12 first-round PCR reactions were performed per replicate, with 18 cycles of amplification (all primer sequences provided in **Table S3**). All first-round PCR reactions from each replicate were then pooled. For the second round of PCR, 5 µL of the pooled first-round PCR product and primers containing barcodes and Illumina sequencing adaptors were used, with 6 to 8 cycles of amplification. PCR products were purified using Agencourt AMPure XP beads (Beckman Coulter Life Sciences). The amplicons were sequenced on an Illumina NextSeq500 with 75 bp single-end reads. sgRNA sequences were extracted from fastq files using an in-house Galaxy script, and normalized read counts were calculated. MAGeCK analysis^48^ was performed to identify genes targeted by depleted sgRNAs at day 15 compared to day 0.

### CRISPRi-mediated gene knockdown

Individual sgRNAs were cloned into pU6-sgRNA lentiviral vector (Addgene #60955) as previously described^77^ (sequences provided in **Table S3)**. CRISPRi cells were transduced with lentiviral vectors in medium containing 6 µg/mL of polybrene (Millipore). After transduction, cells were cultured for 2 days, followed by selection in 2 µg/mL puromycin for 3 days prior to analysis. For *CRNDE* knockdown experiments, *CRNDE* sg1 was used unless otherwise specified, due to its superior knockdown efficiency compared to *CRNDE* sg2.

### Cell proliferation assay

Cells were seeded at a density of 2 ×10^5^ cells per well in 6-well plates in 2 mL of media. Cells were passaged every 3 days and re-seeded at a density of 2 ×10^5^ cells per well after each passage. A Countess cell counter (Thermo Fisher Scientific) was used to quantify cells at each passage, and cumulative cell numbers were plotted. All growth curve experiments were performed with 3 biological replicates.

### RNA isolation and qRT-PCR

Total RNA was extracted from cells using the QIAGEN miRNeasy Mini kit with on-column DNase digestion. cDNA was synthesized from 1 µg of total RNA using the Primescript RT Master Mix (Takara) according to the manufacturer’s instructions. SYBR Green PCR master Mix (Applied Biosystems) was used for qPCR reactions. RNA expression levels were normalized to *GAPDH* using the ΔΔCt method. Primer sequences used for qRT-PCR are provided in **Table S3**.

### Oligo(dT)-primed RT-PCR

To detect *CRNDE* splicing variants by RT-PCR, oligo(dT)-primed reverse transcription was performed using SuperScript IV Reverse Transcriptase (Thermo Fisher Scientific). In brief, 500 ng of total RNA was mixed with 1 μL of 50 μM oligo(dT)20 primer and 1 μL of 10 mM dNTP mix, in a total volume of 13 μL. The mixture was heated at 65°C for 5 minutes and then chilled on ice for 1 minute. For the RT reaction, 4 μL of 5x SSIV buffer, 1 μL of 100 mM DTT, 1 μL of RNase Inhibitor, and 1 μL of SuperScript IV enzyme were added to the annealed RNA mixture. The final reaction was incubated at 50°C for 30 minutes, followed by enzyme inactivation at 80°C for 10 minutes before using cDNA for subsequent PCR reactions. Primer sequences used for RT-PCR are provided in **Table S3**.

### EdU incorporation assay

EdU incorporation assays were performed using the Click-iT Plus EdU Alexa Fluor 488 Imaging Kit (Thermo Fisher Scientific) according to the manufacturer’s instructions. 786-O cells were seeded in 4-well Nunc Lab-Tek II chamber slides (50,000 cells/well, Thermo Fisher Scientific) and cultured for 24 hours. Half of the medium was then replaced with fresh medium containing 20 μM EdU, and cells were incubated for 3 hours. Cells were then fixed with 3.7% formaldehyde and permeabilized with 0.5% Triton X-100. Deoxyuridine incorporation was detected using Click-iT chemistry according to the manufacturer’s instructions. DAPI (2.5 µg/mL) was used for nuclear staining. Cells were imaged using a Zeiss LSM980 confocal microscope. Each experiment was conducted three times, with approximately 200 cells counted per replicate. Images were analyzed using Fiji (ImageJ v1.52p).

### Western blotting

Cells were lysed in 2× NuPAGE LDS Sample Buffer (Thermo Fisher Scientific) for 5 minutes at 95°C. Proteins were separated on 4-12% Bis-Tris NuPAGE gels (Thermo Fisher Scientific) and wet transferred onto nitrocellulose membranes (0.45 μm, Cytiva). Blocking was performed in TBST containing 5% non-fat milk for 1 hour at room temperature. Membranes were incubated with primary antibodies overnight at 4°C, followed by incubation with IR dye-labeled secondary antibodies for 1 hour at RT. Blots were imaged using a LICOR Odyssey imager. All antibodies are listed in **Table S4**.

### Cloning and stable expression of rescue constructs

Tet-On 3G inducible PiggyBac expression vector XLone-GFP (Addgene #96930)^51^ was used for rescue experiments. Rescue constructs were cloned into this vector between the KpnI and SpeI restriction sites using NEBuilder HiFi DNA Assembly Master Mix (NEB) according to the manufacturer’s instructions. Sequences of all rescue constructs are provided in **Table S3**. For constructs that contained the *CRNDE* sgRNA1 sequence, the PAM sequence was mutated for sgRNA resistance. Stable expression of *CRNDE* rescue constructs was achieved by transfecting 786-O cells with XLone-*CRNDE* plasmids along with a plasmid encoding the PiggyBac transposase driven by a CAG promoter, derived from pCMV-hyPBase^81^. Transfected cells were selected with 10 μg/mL blasticidin and single-cell clones were expanded. Rescue constructs were induced using 0.5 μg/mL of doxycycline (Sigma).

### Northern blot analysis

Total RNA (20 μg) or RNA from nucleolar fractions (10 μg) was separated on 1% formaldehyde-agarose gels for larger RNA, or 6% TBE-urea polyacrylamide gels for smaller RNA, and then transferred to BrightStar-Plus nylon membranes (Thermo Fisher Scientific). Membranes were crosslinked using a 254 nm UV crosslinker at 120 mJ/cm^2^. *CRNDE*^UCE^ probes were PCR amplified using primers specific to the UCE and radiolabeled with the Random Primed DNA Labeling Kit (Roche). Probe hybridization was performed using ULTRAhyb (Thermo Fisher Scientific) according to the manufacturer’s instructions. For pre-rRNA northern blots, ITS1 and ITS2 oligo probes were radiolabeled with T4 Polynucleotide Kinase (NEB). Probe hybridization was performed using ULTRAhyb-Oligo (Thermo Fisher Scientific) buffer, following the manufacturer’s instructions. Primers and oligo probe sequences are provided in **Table S3**.

### Measurement of *CRNDE*^UCE^ copy number

The UCE-containing intron of *CRNDE* was in vitro transcribed using the HiScribe T7 High Yield RNA Synthesis Kit (NEB) and used to generate a standard curve with defined numbers of *CRNDE*^UCE^ copies. For each cell line, total RNA was extracted from 1 × 10^6^ cells and reverse transcribed into cDNA for qPCR analysis along with the in vitro transcribed RNA to determine copy number.

### Subcellular fractionation

Fractionation of cytoplasm and nuclei was performed as described previously^82,83^. Briefly, cells were harvested by trypsinization and lysed in 250 µL cytoplasmic lysis buffer (10 mM HEPES pH 7.5, 10 mM KCl, 1.5 mM MgCl_2_, 0.34M sucrose, 0.1% Triton X-100) supplemented with 100 U SUPERase·In RNase Inhibitor (Thermo Fisher Scientific). Following a 10 minutes incubation on ice, the sample was centrifuged at 1300 g for 5 minutes at 4°C. The supernatant, representing the cytoplasmic fraction, was collected and centrifuged at 16,000 g for 15 minutes at 4°C to clear debris. The initial pellet, representing the nuclear fraction, was washed in cytoplasmic lysis buffer to remove residual cytoplasm. The pellet was then resuspended in 250 µL nuclear lysis buffer [50 mM Tris-HCl pH 8, 500 mM NaCl, 1.5 mM MgCl_2_, 0.5% NP-40, and 100 U SUPERase·In RNase Inhibitor (Thermo Fisher Scientific)]. RNA was extracted from each of these fractions using the miRNeasy Mini Kit (Qiagen) with on-column DNase digestion, following the manufacturer’s instructions. Primer sequences used for qRT-PCR are provided in **Table S3**.

### Pre-ribosome sequential extraction

Pre-ribosome sequential extraction (PSE) was performed as described previously^56^. 7 × 10^6^ cells were pelleted and resuspended in 0.5 mL of SN1 buffer (20 mM HEPES pH 7.5, 130 mM KCl, 10 mM MgCl₂, 0.05% Igepal CA-630) supplemented with 1x complete protease inhibitor cocktail (Roche) and 600 U/mL RNasin (Promega). The suspension was centrifuged at 1300 × g for 4 minutes at 4°C. The resulting supernatant was collected and stored as the SN1 fraction. The pellet was washed with 0.5 mL of SN1 buffer, then resuspended in 0.3 mL of SN2 buffer (10 mM HEPES pH 7.5, 10 mM NaCl, 5 mM MgCl₂, 0.1% Igepal CA-630, 0.5 mg/mL heparin,) supplemented with 600 U/mL RNasin (Promega) and 50 U TURBO DNase (Thermo Fisher Scientific). The mixture was incubated for 10 minutes at room temperature with gentle mixing. The lysate was then centrifuged at 12,300 × g for 10 minutes at 4°C, and the supernatant was collected as the SN2 fraction. The remaining pellet was resuspended in 0.4 mL of SN3 buffer [20 mM HEPES pH 7.5, 200 mM NaCl, 4 mM EDTA, 0.1% Igepal CA-630, 0.04% sodium deoxycholate, 4 mM imidazole, 0.1 mg/mL heparin, 1 mM DTT, 1x complete protease inhibitor cocktail (Roche), 600 U/mL RNasin (Promega)]. This mixture was incubated for 20 minutes at room temperature with gentle rotation. The extract was centrifuged at 12,300 × g for 10 minutes at 4°C, and the supernatant was collected as the SN3 fraction. For western blotting, samples from equal cell equivalents of the SN1, SN2, and SN3 fractions (in a volumetric ratio of 10:6:8) were mixed with 2× NuPAGE LDS Sample Buffer (Thermo Fisher Scientific). RNA was extracted from each fraction using the RNeasy Mini Kit (Qiagen). All antibodies are listed in **Table S4**. Primer sequences used for qRT-PCR are provided in **Table S3**.

### RNA fluorescence in situ hybridization (RNA FISH)

RNA FISH was performed as previously described^84,85^. Briefly, DIG-labeled anti-sense RNA probes for *CRNDE*^UCE^ were synthesized by in vitro transcription using a DIG-labeling mix (Roche) and purified using Micro Bio-Spin P-30 chromatography columns (Bio-Rad). Primers used for amplification of DNA template are provided in **Table S3**. 786-O cells (5 × 10^4^) were seeded in 4-well Nunc Lab-Tek II chamber slides (Thermo Fisher Scientific) and cultured for 2 days. Cells were rinsed twice in DEPC-treated phosphate buffered saline (PBS) and fixed in 4% paraformaldehyde for 10 minutes at room temperature. Cells were then washed once with DEPC-treated PBS and permeabilized with 0.5% Triton X-100 for 10 minutes at room temperature. After permeabilization, cells were washed twice with DEPC-treated PBS and incubated with pre-hybridization buffer (50% formamide, 2X SSC, 1X Denhardt’s solution, 10 mM EDTA, 0.1 mg/mL yeast tRNA, 0.01% Tween-20) for 1 hour. DIG-labelled RNA probes (10 ng/μL) were diluted in hybridization buffer (prehybridization buffer with 5% dextran sulfate), denatured at 75 °C for 10 minutes, and hybridized at 55 °C for 18 hours. Following hybridization, samples were washed, treated with RNase A, and blocked for 1 hour at room temperature with blocking reagent (Roche). Detection of DIG-labeled probes were performed by incubation with a mouse monoclonal anti-DIG primary antibody (Roche), followed by incubation with Cy3-labeled goat anti-mouse IgG secondary antibody (EMD Millipore). Nuclei were stained with DAPI (2.5 µg/mL) for 1 minute, washed with TBST, and coverslips were mounted with SlowFade Diamond Antifade Mountant with DAPI (Thermo Fisher Scientific). Images were acquired using a Zeiss LSM980 confocal microscope with a 63x oil objective. Images were analyzed using Fiji (ImageJ v1.52p).

### Sucrose gradient ultracentrifugation and fractionation

Polysome fractionation by sucrose gradient ultracentrifugation was performed as described^86^ with minor modifications to the protocol. Cells were incubated with 100 µg/mL cycloheximide (Sigma) in growth medium for 10 minutes, then washed twice with ice-cold 1x PBS containing 100 µg/mL cycloheximide. Cells were scraped in 5 mL of ice-cold 1x PBS containing 100 µg/mL cycloheximide and collected by centrifugation. Cell pellets were lysed in 500 µL hypotonic buffer (5 mM Tris-HCl pH 7.5, 2.5 mM MgCl_2_, 1.5 mM KCl, 100 µg/mL cycloheximide, 2 mM DTT, 0.5% Triton X-100, and 0.5% sodium deoxycholate) supplemented with 1x EDTA-free protease inhibitor cocktail (Roche) and 200 U/mL RNasin (Promega). Lysates were centrifuged at 16,000 x g for 7 minutes at 4° C. Supernatants were collected, diluted with lysis buffer as needed so all samples had an equivalent OD_260_. The samples were then loaded onto a 5–50% sucrose gradient. Gradients were ultracentrifuged at 36,000 rpm for 2 hours at 4°C using a TH-641 rotor (Thermo Fisher Scientific) and fractionated on a Piston Fractionator (BioComp).

### RNA antisense purification and mass spectrometry (RAP-MS)

RAP-MS coupled with organic phase separation was performed as previously described^58,60^ with the following modifications:

#### 1. UV crosslinking

786-O cells were grown to approximately 80-90% confluence in 15 cm dishes. The cells were washed twice with ice-cold PBS and 6 mL ice-cold PBS was added to each plate. Crosslinking was performed at 254 nm UV with an energy setting of 400 mJ/cm² using a Spectrolinker XL-1500 (Spectronics). The cells were then scraped and pelleted by centrifugation at 1000 x g for 5 minutes.

#### 2. Organic Phase Separation

20 million cells were lysed in 900 µL of total cell lysis buffer [10 mM Tris-HCl pH 7.5, 500 mM LiCl, 0.5% Triton X-100, 0.2% SDS, 0.1% sodium deoxycholate, 400 U SUPERase·In RNase Inhibitor (Thermo Fisher Scientific), 1x EDTA-free protease inhibitor cocktail (Roche)]. Samples were incubated on ice for 10 minutes, during which the cell lysate was passed 3-5 times through a 26-gauge needle attached to a 1 mL syringe. The samples were then sonicated using a Bioruptor Plus (30 seconds on/30 seconds off, 5 cycles). Lysates were treated with 20 U TURBO DNase (Thermo Fisher Scientific) for 30 minutes at 37°C. Each sample was split into 4 tubes and 750 µL TRIzol LS was added to 250 µL of lysate, followed by the addition of 200 µL of chloroform per 1 mL of TRIzol mix. Samples were centrifugated at 20,000 x g for 15 minutes. The aqueous phase was removed, and the interphases from each sample were collected in a 2 mL tube. The interphase was gently washed twice with 1 mL of low SDS buffer (50 mM Tris-HCl, 1 mM EDTA, 0.1% SDS) by centrifugation at 5000 g for 2 minutes at room temperature, and the supernatant was discarded. After washing, the flakes were disintegrated by pipetting into 100 µL of low SDS buffer and 900 µL of ethanol. The samples were incubated at -20°C for at least 2 hours. The samples were then centrifuged at 20,000 x g for 30 minutes at 4°C and the pellet was washed with 70% ethanol. The pellet was then resuspended in 500 µL of high SDS buffer (20 mM Tris-HCl pH 7.5, 1 mM EDTA, 0.4% SDS) and incubated for 20 minutes at 56°C. The samples were then centrifuged at 1000 x g for 2 minutes to separate the soluble from the insoluble material. The supernatant was collected and the volume was adjusted to 1 mL using concentrated RAP-MS hybridization buffer (final concentrations: 10 mM Tris-HCl pH 7.5, 500 mM LiCl, 5 mM EDTA, 0.2% SDS, 0.5% DDM, 4 M urea, 2.5 mM TCEP) with 400 U SUPERase·In RNase Inhibitor (Thermo Fisher Scientific) and 1x EDTA-free protease inhibitor cocktail (Roche).

#### 3. RAP-MS

To pull-down *CRNDE*^UCE^, 60-nucleotide 5’ biotinylated ASOs (IDT) were used. 200 pmol ASOs per 20 million cells were prepared by heating ASOs at 85°C for 3 minutes, mixing with streptavidin magnetic beads (NEB), incubating for 1 hour at room temperature, and then adding to 1 mL sample in hybridization buffer prepared in step 2. Hybridization was carried out at 60°C for 2 hours. Post-hybridization, the beads were washed, and elution was performed with water at 70°C for 5 minutes. Label-free semi-quantitative mass spectrometry was performed at the UT Southwestern Proteomics Core using reverse-phase LC-MS/MS and an Orbitrap Fusion Lumos mass spectrometer. Raw MS data files were analyzed using Proteome Discoverer v2.4 SP1 (Thermo Fisher Scientific), with peptide identification performed using Sequest HT searching against the human protein database from UniProt.

To directly detect protein interactors by western blotting, proteins were eluted from streptavidin-coated magnetic beads after the final wash by resuspending beads in NuPAGE LDS Sample Buffer (Thermo Fisher Scientific) and incubating for 10 minutes at 70° C. Eluted proteins were detected by western blotting as described above. All antibodies are listed in **Table S4**.

### UV crosslinking and RNA immunoprecipitation (UV-RIP)

UV-RIP was performed as previously described with the following modifications^85^. 786-O cells (1.5 × 10^7^) were washed with ice-cold PBS and UV crosslinked on ice at 254 nm with 400 mJ/cm² using a Spectrolinker XL-1500 (Spectronics). Cells were scraped in cold PBS and pelleted by centrifugation. Cell pellets were lysed in 1 mL of cold lysis buffer [50 mM Tris-HCl pH 7.5, 100 mM NaCl, 1% NP-40, 0.1% SDS, 0.5% sodium deoxycholate, 1x complete protease inhibitor cocktail (Roche), 400 U/mL RNasin (Promega)] for 30 minutes on ice followed by sonication using a Bioruptor Plus (15 seconds on/60 seconds off, 5 cycles). Lysates were cleared by centrifugation at 16,000 x g for 15 minutes at 4°C. 50 µL (1.5 mg) of pre-washed Protein G Dynabeads (Thermo Fisher Scientific) coupled with 5 ug of antibody was added to the cleared lysate and rotated for 2 hours at 4° C. Beads were washed three times with 900 µL cold High Salt Wash Buffer (50 mM Tris-HCl pH 7.5, 1M NaCl, 1 mM EDTA, 1% NP-40, 0.1% SDS, 0.5% sodium deoxycholate) and then washed three times with 500 µL wash buffer (20 mM Tris-HCl pH 7.5, 10 mM MgCl_2_, 0.2% Tween-20). After the final wash, each sample was resuspended in 100 µL of wash buffer. 30 µL of the sample was used for western blotting. The remainder was treated with proteinase K (NEB) followed by RNA extraction with TRIzol.

### Expression and purification of eIF6-3XFlag

eIF6-3XFlag was cloned into pLenti-CMV-Blast (Addgene #17486)^87^ using the Esp3I and BamH1 restriction sites. After transducing HEK293T cells with this lentiviral construct, cells were cultured for 2 days, followed by selection in 10 µg/mL blasticidin for 10 days. HEK293T eIF6-3XFlag cells (1 × 10^7^ cells) were harvested and resuspended in 500 µL of lysis buffer [50 mM Tris-HCl pH 7.5, 300 mM NaCl, 0.5% NP-40, 0.1% sodium deoxycholate, 5 mM MgCl_2_, 2 mM DTT, 1x complete protease inhibitor cocktail (Roche)] for 30 minutes on ice, followed by sonication using a Bioruptor Plus (15 seconds on/60 seconds off, 3 cycles). An equal volume of dilution buffer [50 mM Tris-HCl pH 7.5, 0.5% NP-40, 2 mM DTT, 1x complete protease inhibitor cocktail (Roche)] was then added. The samples were cleared by spinning at 16,000 x g for 15 minutes. 25 µL of anti-Flag magnetic beads (Thermo Fisher Scientific) were added per approximately 1 mg of total protein in the lysate, and the samples were incubated at 4°C for 4 hours. The beads were washed four times for 5 minutes each with wash buffer (50 mM Tris-HCl pH 7.5, 300 mM NaCl, 0.05% sodium deoxycholate, 0.5% NP-40). eIF6 was eluted from the beads in 100 µL of elution buffer (50 mM Tris-HCl pH 7.5, 150 mM NaCl, 1 mM EDTA, 0.05% NP-40, 10% glycerol, 1× EDTA-free Protease Inhibitor Cocktail, 1.5 mg/mL 3×Flag peptide) at 16°C for 30 minutes at 800 rpm. The elution step was repeated and eluates were combined.

### Electrophoretic mobility shift assay (EMSA)

EMSAs were performed using purified eIF6-3XFlag protein and RNA synthesized by in vitro transcription with the HiScribe T7 High Yield RNA Synthesis Kit (NEB). Primers used for amplification of UCE intron or UCE intron fragment templates are provided in **Table S3**. In vitro transcribed RNAs were folded by heating at 65°C for 5 minutes, then slowly cooled down to room temperature. Protein-RNA binding was carried out with purified eIF6-3XFlag (200 nM) and folded RNA (0.5 pmol) in binding buffer (10 mM HEPES pH 7.5, 3 mM MgCl_2_, 40 mM KCl, 5% glycerol, and 1 mM DTT). After incubation at room temperature for 30 minutes, samples were immediately loaded onto a 1.2% agarose gel and electrophoresis was carried out as previously described^88^. RNAs were detected using 2x SYBR Gold (Thermo Fisher Scientific) staining.

### Immunolocalization of eIF6

786-O cells (5 × 10^4^) were seeded in 4-well Nunc Lab-Tek II chamber slides (Thermo Fisher Scientific) and cultured for 2 days. Cells were rinsed twice with PBS and fixed with 4% paraformaldehyde for 15 minutes at room temperature. Cells were then washed twice with PBS. Permeabilization was performed with 0.5% Triton X-100 in PBS for 10 minutes at room temperature, followed by two washes with PBS. Samples were washed with 1X PBST (PBS + 0.1% Tween 20) for 5 minutes. Blocking was performed with 3% BSA in PBST for 30 minutes at room temperature. After blocking, samples were incubated with primary anti-eIF6 antibody diluted 1:100 in 1% BSA in PBST overnight at 4°C. Samples were then washed three times with 1X PBST. Secondary antibody incubation was performed with goat anti-mouse IgG cross-adsorbed secondary antibody (Thermo Fisher Scientific) diluted 1:200 in 1% BSA in PBST for 1 hour at room temperature, followed by three washes with PBS. Nuclei were stained with DAPI (2.5 µg/mL) for 1 minute, washed with PBS, and then coverslips were mounted with Immumount (Thermo Fisher Scientific). Images were acquired using a Zeiss LSM980 confocal microscope with a 63x oil objective. Images were analyzed using Fiji (ImageJ v1.52p).

### Sucrose gradient fractionation of pre-ribosomes

Pre-ribosomes were analyzed by sucrose gradient ultracentrifugation as described previously^56^. Freshly prepared 400 µL samples of SN3 fraction were loaded onto a 7–50% sucrose gradient. The gradients were prepared in a buffer containing 20 mM HEPES pH 7.5, 200 mM NaCl, 4 mM EDTA, 0.1 mg/mL heparin, and 1 mM DTT. Ultracentrifugation was performed at 39,000 rpm for 165 minutes at 4°C using a TH-641 rotor (Thermo Fisher Scientific). The samples were then fractionated using a Piston Fractionator (BioComp) to evaluate pre-ribosome profiles. To detect eIF6 distribution, 0.5 mL fractions were collected from the gradient. From each fraction, 380 µL was transferred to a corresponding tube containing 130 µL of 50% trichloroacetic acid (TCA). The samples were mixed thoroughly and stored at -20°C overnight to precipitate the proteins. The proteins in each sample were pelleted by centrifugation at 16,000 x g for 15 minutes at 4°C. The pellets were then washed twice with 200 µL of 100% acetone and dried using a speed vacuum apparatus for 10 minutes. Each pellet was resuspended in 30 µL of 2x LDS Sample Buffer and neutralized with 15 µL of 2 M Tris-HCl pH 8 for western blot analysis.

### Co-immunoprecipitation

786-O cells (5 × 10^6^) were harvested and lysed in 500 μL IP buffer [20 mM Tris-HCl pH 7.5, 150 mM NaCl, 1 mM EDTA, 0.5% NP-40, 1x complete protease inhibitor cocktail (Roche)] for 30 minutes on ice, followed by sonication using a Bioruptor Plus (10 seconds on/60 seconds off, 3 cycles). Lysates were centrifuged at 16,000 x g for 15 minutes, and supernatant was collected. The cleared lysates were incubated with 2 μg anti-eIF6 or anti-NMD3 antibodies overnight at 4°C with rotation. Then, 30 μL Protein G Dynabeads (Thermo Fisher Scientific) were added to the mixture and incubated for 2 hours at 4°C, followed by washing four times with IP buffer. To elute the protein complex, 2x NuPAGE LDS Sample Buffer (Thermo Fisher Scientific) was added to the samples, which were then boiled for 5 minutes. The eluted proteins were analyzed by western blotting. All antibodies are listed in **Table S4**.

### RPL23a-SNAP labeling

RPL23a-SNAP sequences (IDT) were cloned into a previously generated AAVS1 donor plasmid^85^ using the XhoI and MfeI restriction sites. The *AAVS1*/ RPL23a-SNAP knock-in construct was then inserted into the *AAVS1* locus of 786-O CRISPRi cells, as previously described^85,89^. Transfected cells were selected with 500 µg/mL G418 for at least 10 days before introducing sgRNAs for CRISPRi-mediated knockdown. Detection of newly-synthesized RPL23a using SNAP staining was performed as previously described with minor modifications^90,62^. Cells were incubated with SNAP-Cell Block (NEB) at a final concentration of 10 µM in growth medium at 37°C for 30 minutes to quench pre-existing ribosomes. Cells were then washed three times with fresh medium, followed by a 6-8 hour incubation in fresh medium. Cells were then incubated with 2 µM SNAP-Cell Oregon Green (NEB) for 20 minutes at 37°C to label newly synthesized RPL23a. Images were acquired using a Zeiss LSM980 confocal microscope with a 40x objective. Images were analyzed using Fiji (ImageJ v1.52p).

### Detection of RNA:RNA interactions by AMT crosslinking

AMT crosslinking and detection of RNA:RNA interactions were performed as previously described with modifications^64^. 786-O cells (2 × 10^7^) were harvested and resuspended in 4 mL of ice-cold 0.5 mg/mL AMT solution (+AMT) or ice-cold PBS alone (-AMT control) and incubated on ice in the dark for 15 minutes. The samples were then transferred to pre-chilled 10-cm tissue culture dishes, placed on ice under a long-wave UV bulb (350 nm) in a Spectrolinker XL-1500 (Spectronics) approximately 4 cm from the light source, and covered with a 0.3 mm glass plate. Cells were exposed to UV light for 15 minutes (450 seconds twice), then transferred to pre-chilled tubes and spun at 1000 x g for 5 minutes. The cell pellets were resuspended in 500 µL of cell lysis buffer (10 mM Tris-HCl pH 7.5, 500 mM LiCl, 0.5% Triton X-100, 0.2% SDS, 0.1% sodium deoxycholate) containing 200 U SUPERase·In RNase Inhibitor (Thermo Fisher Scientific). Lysates were treated with 20 U TURBO DNase (Thermo Fisher Scientific) for 15 minutes at 37°C, followed by treatment with 0.25 mg/mL proteinase K (NEB) for 30 minutes at 37°C. RNA was extracted using the RNeasy Mini Kit (Qiagen). Input RNA (2 µg) and biotinylated ASO probe (50 pmol) were denatured separately in 5 µL water at 85°C for 3 minutes, then mixed in 300 µL of LiCl Hybridization Buffer (10 mM Tris-HCl pH 7.5, 1 mM EDTA, 500 mM LiCl, 1% Triton X-100, 0.2% SDS, 0.1% sodium deoxycholate, 4 M urea) preheated to 55°C. The hybridization reaction was incubated for three hours at 55°C with shaking at 1,000 rpm. 50 µL streptavidin C1 magnetic beads (Thermo Fisher Scientific) were added and reactions were incubated at 37°C for 45 minutes to capture the complexes. The beads were washed three times with 250 µL of Low Stringency Wash Buffer (1× SSPE, 0.1% SDS, 1% NP-40, 4 M urea) and three times with 250 µL of High Stringency Wash Buffer (0.1× SSPE, 0.1% SDS, 1% NP-40, 4 M urea), each wash for 5 minutes at 60°C. RNA was eluted using RNase H, purified using TRIzol and RNA Clean & Concentrator-5 columns (Zymo Research), and then subjected to qRT-PCR as described above. Primer sequences used for qRT-PCR provided in **Table S3**.

### Quantification and statistical analysis

All experiments were repeated with a minimum of three independent biological replicates as indicated in each figure legend. All values are reported as mean ± SD in each figure. Statistical significance was analyzed using Prism 10 (GraphPad Software). Statistical analysis of CRISPRi screen was performed using MAGeCK^48^.

