## Supplemental Figures for "An ultraconserved snoRNA-like element in long noncoding RNA *CRNDE* promotes ribosome biogenesis and cell proliferation"

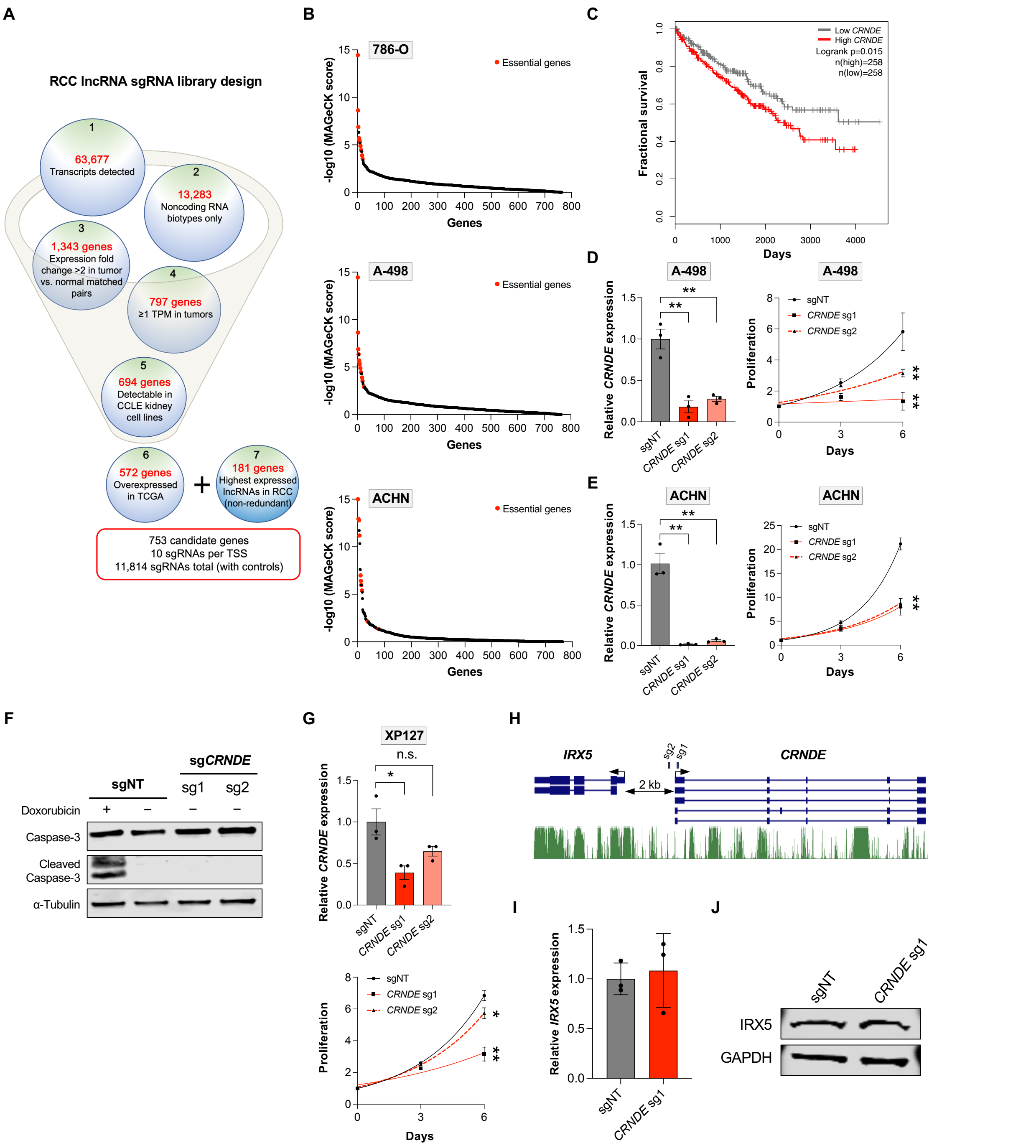


**Figure S1. *CRNDE* is required for RCC cell proliferation, related to Figures 1 and 2**
(**A**) Design of sgRNA library targeting 753 highly expressed lncRNAs in RCC. (**B**) Essential genes labeled in red (*PCNA, RPA1, RPA3, RPS15A, RPS19, RPS11, RPS21, RPL32, RPS18, RPL7A, RPL8, RPL6*) were strongly depleted in CRISPRi screens in all RCC cell lines. (**C**) High expression of *CRNDE* is associated with reduced survival of RCC patients. Analysis performed using GEPIA2[75]. (**D-E**) qRT-PCR analysis of *CRNDE* expression relative to *GAPDH* in A-498 (D) or ACHN (E) CRISPRi cells after lentiviral expression of non-target (sgNT) or *CRNDE*-targeting sgRNAs (left panels). Proliferation was measured by cell counts at the indicated timepoints after plating (right panels). (**F**) Western blot analysis of apoptosis marker cleaved caspase-3 in 786-O CRISPRi cells after lentiviral expression of non-target (sgNT) or *CRNDE*-targeting sgRNAs. Cells treated with 1 µM doxorubicin served as a positive control. (**G**) qRT-PCR analysis of *CRNDE* expression relative to *GAPDH* in patient-derived primary RCC CRISPRi cells after lentiviral expression of non-target (sgNT) or *CRNDE*-targeting sgRNAs (upper). Proliferation was measured by cell counts at the indicated timepoints after plating (lower). (**H**) UCSC genome browser RefSeq and vertebrate conservation (PhastCons) tracks (hg38) showing the *CRNDE* and nearby *IRX5* transcription units. Positions of *CRNDE* sg1 and sg2 indicated. (**I-J**) qRT-PCR analysis of *IRX5* expression relative to *GAPDH* (I) and western blot analysis of IRX5 (J) in 786-O CRISPRi cells expressing the indicated sgRNAs.
Data are represented as mean ± SD (n=3 biological replicates). n.s., not significant; *p<0.05, **p<0.01, calculated by two-tailed t-test.

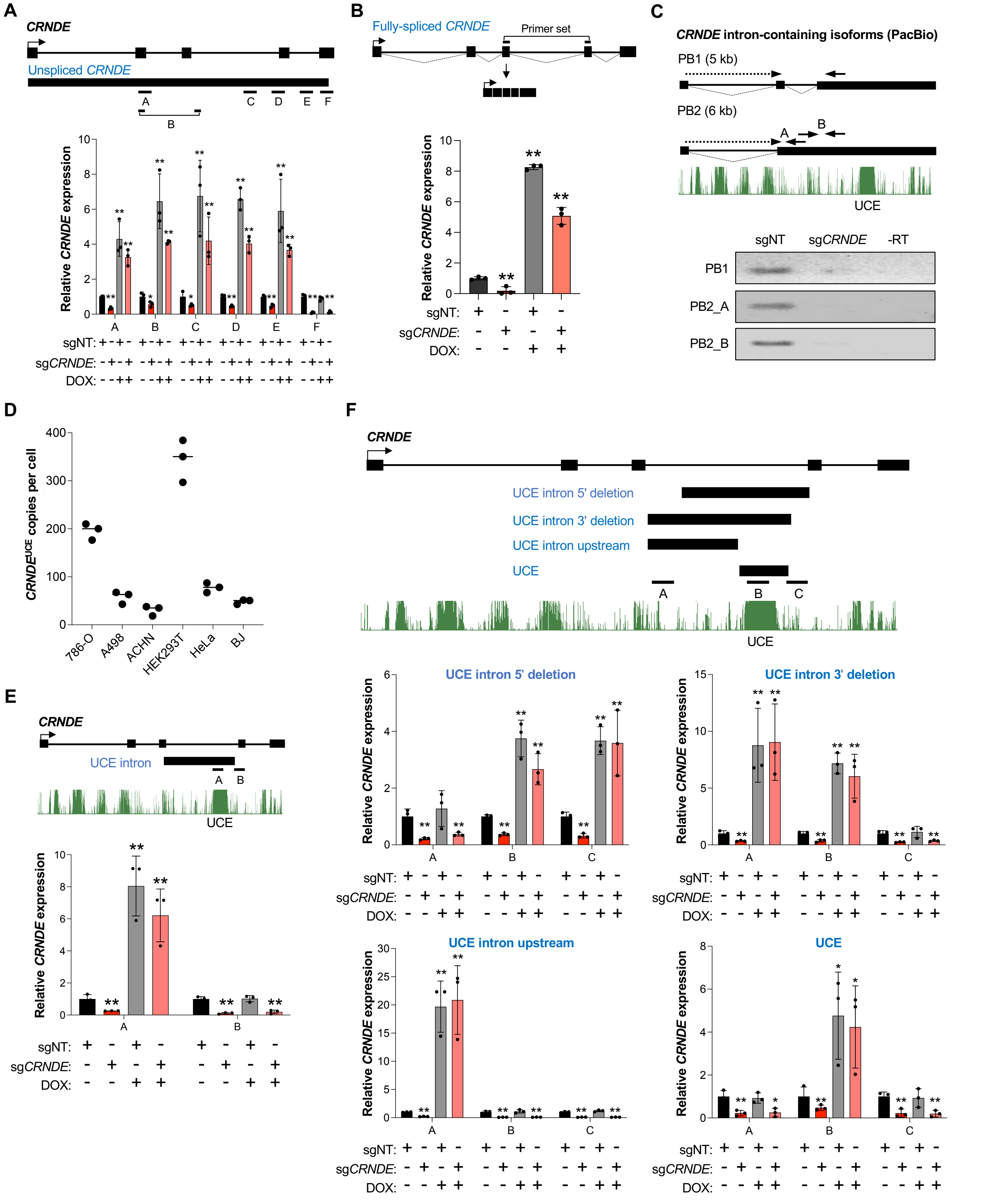


**Figure S2. An ultraconserved element-containing isoform of *CRNDE* promotes RCC proliferation in *trans*, related to Figure 2**

(**A,B,E,F**) qRT-PCR analysis of 786-O CRISPRi cells expressing the indicated dox-inducible *CRNDE* rescue constructs after lentiviral expression of non-target sgRNA (sgNT) or sgRNA targeting endogenous *CRNDE*. The positions of qRT-PCR amplicons are indicated in the transcript schematics. Transcript abundance was normalized to *GAPDH*. (**C**) Upper, schematic representation of intron-containing *CRNDE* isoforms detected by PacBio sequencing (PB1 and PB2), aligned with UCSC genome browser vertebrate conservation track (PhastCons, hg38). Arrows indicate the approximate positions of RT-PCR primers. Bottom, agarose gel image of RT-PCR of oligo-dT primed cDNA from 786-O cells expressing the indicated sgRNAs. (**D**) *CRNDE*$^{UCE}$ transcript copy number per cell in the indicated cell lines, measured by qRT-PCR using RNA from a defined number of cells and a standard curve (see Materials and Methods).
Data are represented as mean ± SD (n=3 biological replicates). *p<0.05, **p<0.01, calculated by two-tailed t-test.

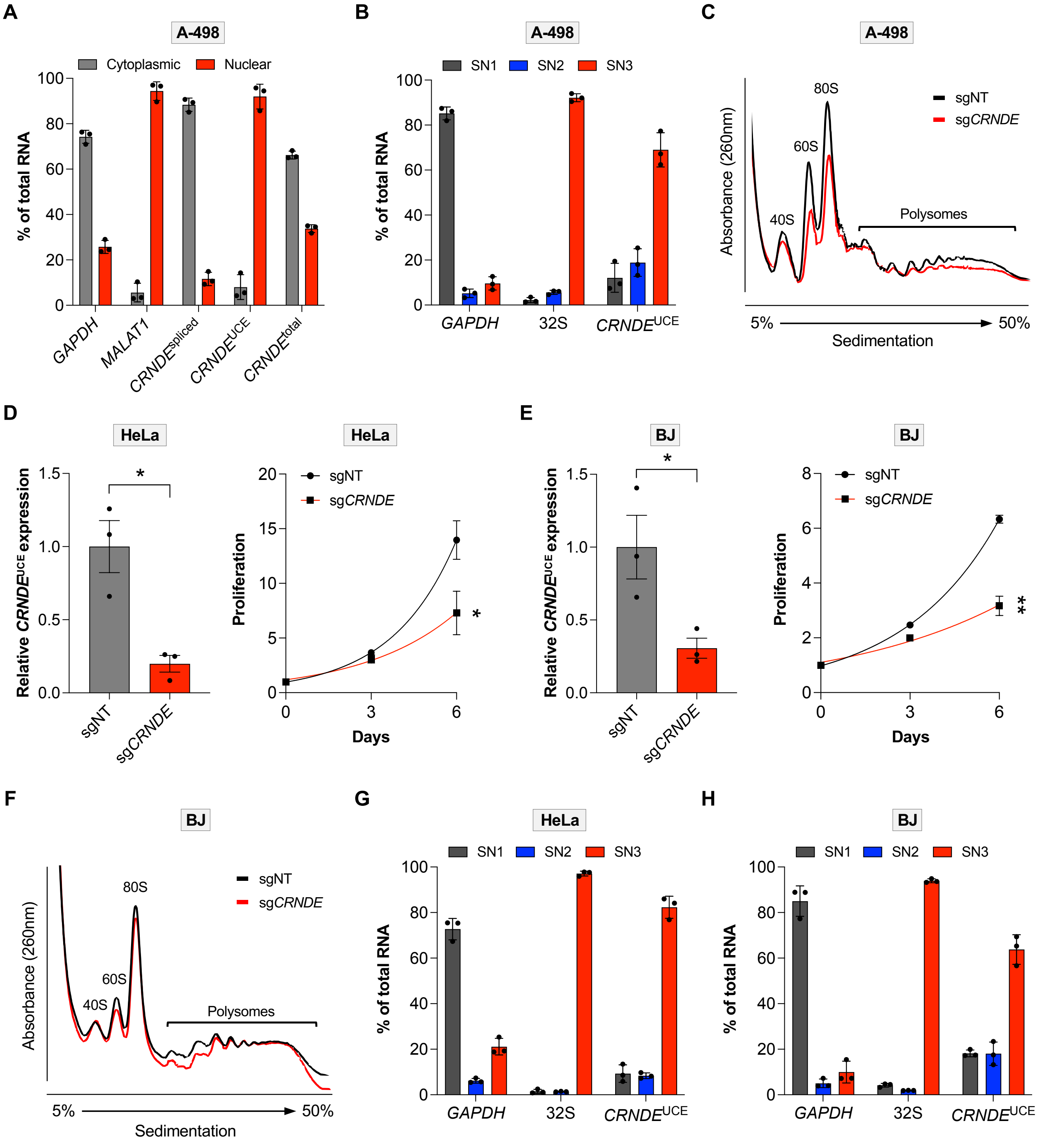


**Figure S3. *CRNDE*$^{UCE}$ localizes to the nucleolus and promotes 60S ribosomal subunit biogenesis, related to Figure 3**

(**A**) Subcellular fractionation and qRT-PCR analysis of A-498 cells using primers that detect fully-spliced *CRNDE* (*CRNDE*$^{spliced}$), *CRNDE* UCE-containing isoforms (*CRNDE*$^{UCE}$), or all *CRNDE* isoforms (*CRNDE*$^{total}$). *GAPDH* and *MALAT1* represent cytoplasmic and nuclear controls, respectively. (**B**) qRT-PCR analysis of *CRNDE*$^{UCE}$ in nucleolar fractions from A-498 cells. *GAPDH* and 32S pre-rRNA represent cytoplasmic and early nucleolar markers, respectively. (**C**) Sucrose gradient fractionation of lysates from A-498 CRISPRi cells after lentiviral expression of non-target (sgNT) or *CRNDE*-targeting sgRNAs. (**D-E**) qRT-PCR analysis of *CRNDE* expression relative to *GAPDH* in HeLa (D) or BJ (E) CRISPRi cells after lentiviral expression of non-target (sgNT) or *CRNDE*-targeting sgRNAs (left panels). Proliferation was measured by cell counts at the indicated timepoints after plating (right panels). (**F**) Sucrose gradient fractionation of lysates from BJ CRISPRi cells after lentiviral expression of non-target (sgNT) or *CRNDE*-targeting sgRNAs. (**G-H**) qRT-PCR analysis of *GAPDH*, 32S pre-rRNA, and *CRNDE*$^{UCE}$ in nucleolar fractions from HeLa cells (G) and BJ fibroblasts (H).

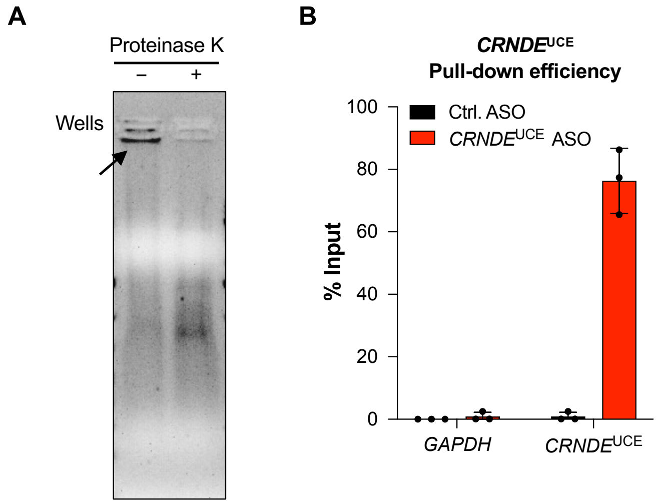


C

| Protein | Abundance | Biological process |
|---|---|---|
| **eIF6** | **3.85E+05** | **Large ribosomal subunit biogenesis** |
| PGM2 | 1.30E+05 | Glycogen catabolic process |
| SLURP1 | 1.04E+05 | Acetylcholine receptor signaling |
| EMD | 1.01E+05 | Nuclear matrix anchoring at nuclear membrane |
| CST1 | 9.61E+04 | Negative regulation of cysteine-type endopeptidase activity |
| LIG1 | 5.64E+04 | Base-excision repair, gap-filling |
| DDX56 | 5.40E+04 | Large ribosomal subunit biogenesis |
| TENM1 | 5.37E+04 | Adenylate cyclase-activating G protein-coupled receptor signaling |
| ANXA8L1 | 3.86E+04 | Negative regulation of phospholipase A2 activity |
| RAB14 | 3.62E+04 | Post-golgi vesicle-mediated transport |
| ELOVL1 | 2.76E+04 | Fatty acid elongation |
| FN1 | 2.35E+04 | Cell-cell adhesion mediated by integrin |

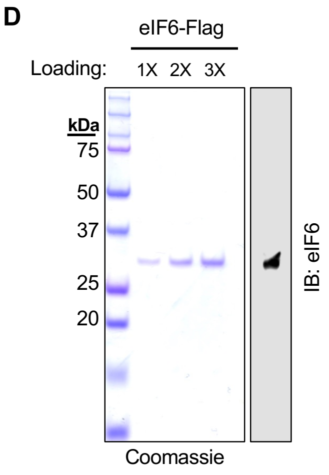


**Figure S4. *CRNDE*$^{UCE}$ directly interacts with eIF6, related to Figure 4**
(**A**) Enrichment of crosslinked RNPs by organic phase separation was confirmed by agarose gel electrophoresis and ethidium bromide staining. Crosslinked RNPs are trapped in the wells and RNA is released by proteinase K digestion. (**B**) qRT-PCR analysis of *CRNDE*$^{UCE}$ recovery after pull-down with ASOs. *GAPDH* served as a negative control. (**C**) Proteins exclusively detected in *CRNDE*$^{UCE}$ ASO pull-downs, but absent in scrambled ASO pull-downs, ranked by detected abundance. (**D**) Coomassie stain and western blot of purified eIF6 used for in vitro binding experiments.

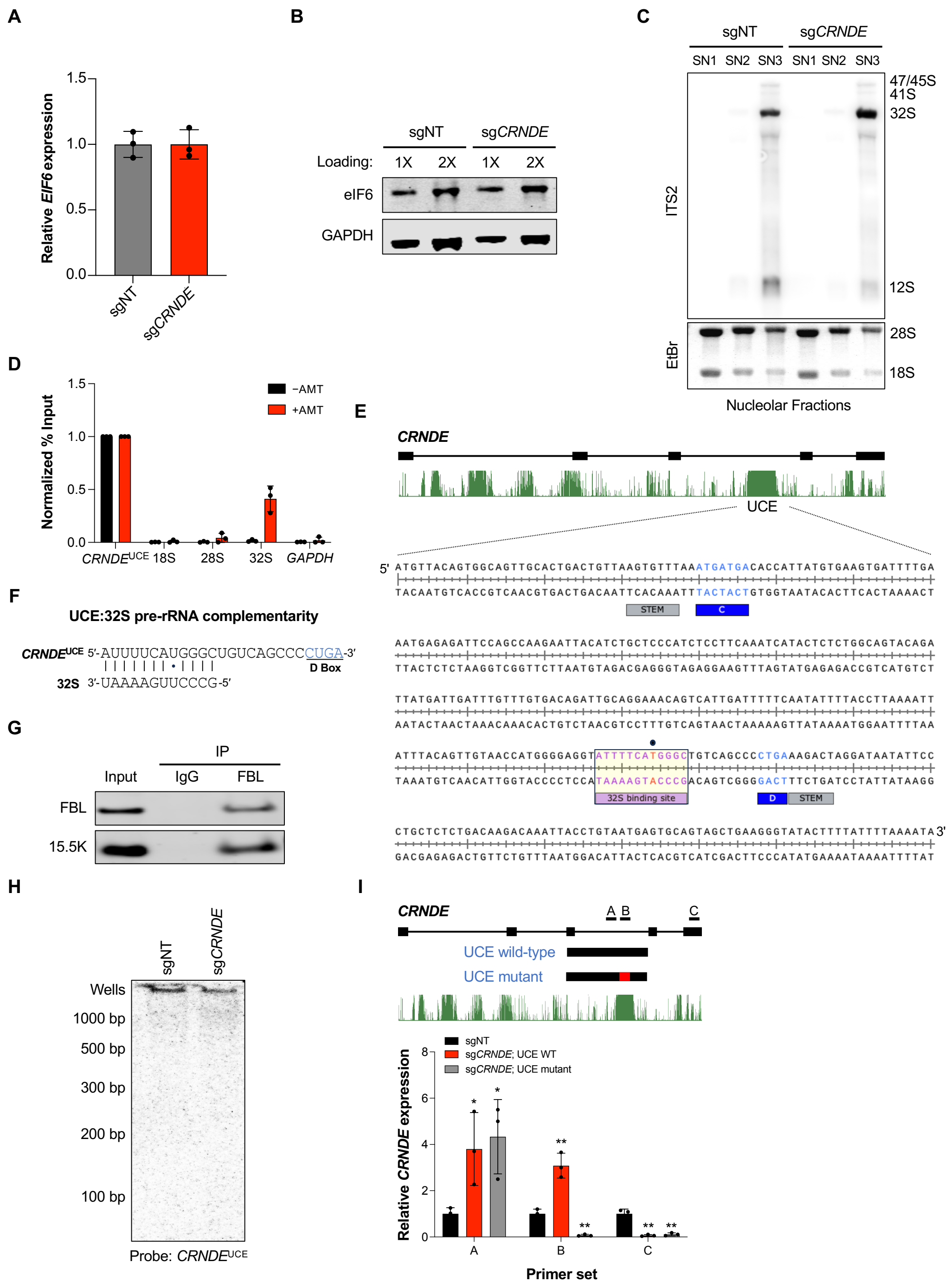


**Figure S5. The *CRNDE* UCE is a C/D box snoRNA-like element that base-pairs with 32S pre-rRNA and promotes eIF6 loading, related to Figures 5 and 6**

(**A-B**) qRT-PCR analysis of *EIF6* expression relative to *GAPDH* (A) and western blot analysis of eIF6 (B) in 786-O CRISPRi cells expressing the indicated sgRNAs. (**C**) Northern blot analysis of rRNA precursors with ITS2 probe in nucleolar fractions from 786-O CRISPRi cells expressing the indicated sgRNAs. EtBr, ethidium bromide. (**D**) qRT-PCR analysis of the indicated transcripts after AMT crosslinking and pull-down of *CRNDE*$^{UCE}$ with ASOs under denaturing conditions. Omission of AMT prior to UV exposure served as a negative control for this experiment. Enrichment was normalized to input. (**E**) The complete genomic sequence of the *CRNDE* UCE with C box, D box, terminal stem, and rRNA complementarity site indicated. (**F**) Predicted base-pairing between UCE and 32S pre-rRNA. (**G**) Western blot analysis of FBL or IgG immunoprecipitates from 786-O cells. Co-immunoprecipitation of 15.5K, another C/D box snoRNP component, was observed. RNA from this experiment was analyzed in Figure 6E. (**H**) Northern blot analysis of total RNA from 786-O CRISPRi cells using a probe complementary to the UCE failed to detect any small RNA species. (**I**) qRT-PCR analysis of 786-O CRISPRi cells expressing dox-inducible *CRNDE* UCE intron rescue constructs with or without mutations in the putative pre-rRNA binding site after lentiviral expression of non-target sgRNA (sgNT) or sgRNA targeting endogenous *CRNDE*. The positions of qRT-PCR amplicons are indicated in the transcript schematic. Transcript abundance was normalized to *GAPDH*.
Data are represented as mean ± SD (n=3 biological replicates). *p<0.05, **p<0.01, calculated by two-tailed t-test.

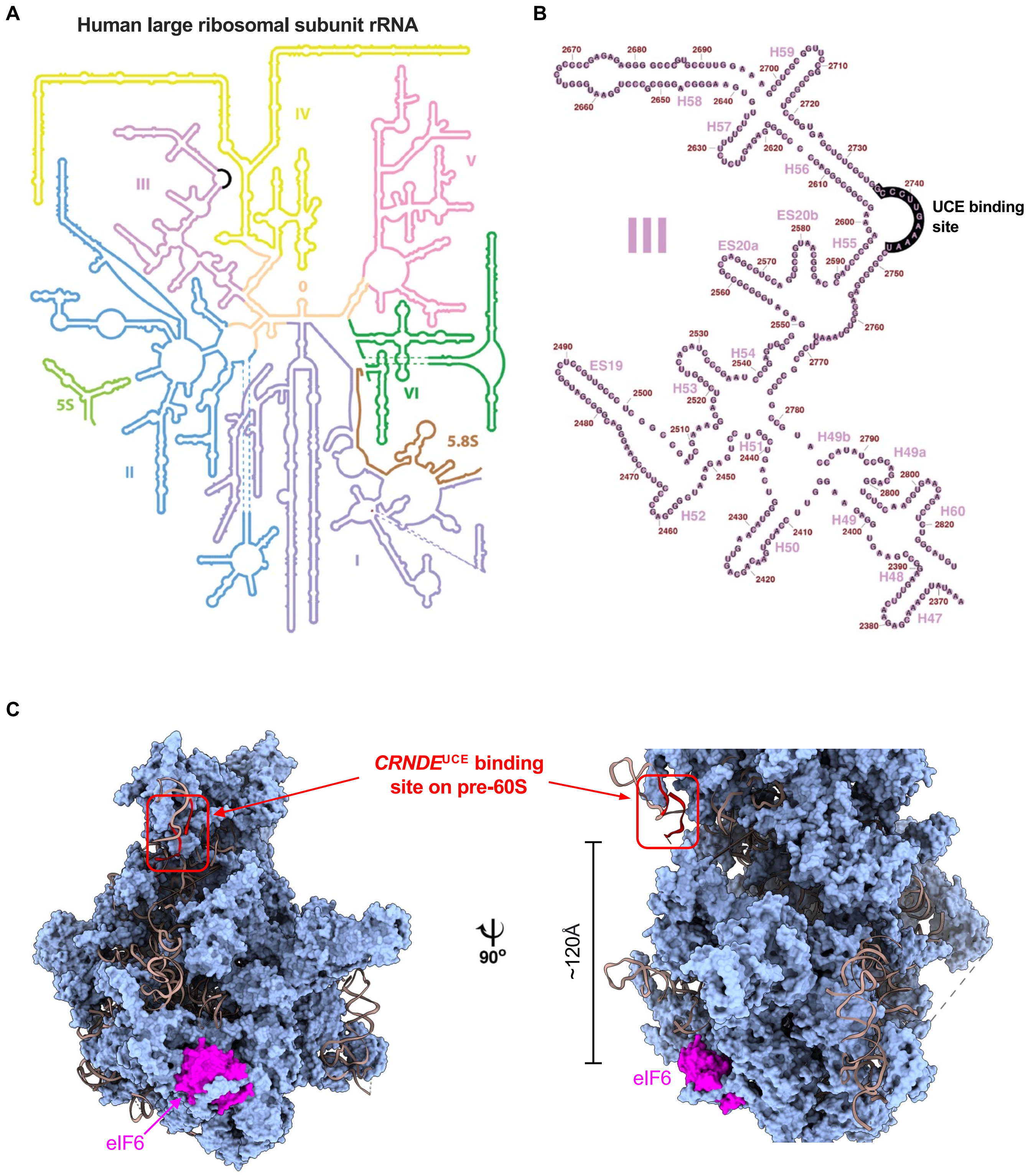


**Figure S6. Binding site for the *CRNDE* UCE in rRNA and pre-60S particle, related to Figure 6**
(**A**) Secondary structure map of human rRNA in 60S subunit, adapted from Petrov et al.[76] Black segment represents UCE binding site. (**B**) Secondary structure map of human 28S rRNA domain III, showing UCE binding site in the linker between helices H55 and H56. (**C**) Cryo-EM structure of a human nucleolar pre-60S particle containing eIF6 (PDB: 8FKV)[67]. The sequence in 32S pre-rRNA predicted to base-pair with the *CRNDE* UCE is located in a structurally unresolved loop on the particle surface (red strands indicate proximal and distal segments of unresolved sequence; brown strands represent remainder of 32S pre-rRNA). eIF6 (magenta) binds to the same face of the particle, approximately 120Å away.
